# Single Cell Profiling Reveals Sex, Lineage and Regional Diversity in the Mouse Kidney

**DOI:** 10.1101/673335

**Authors:** Andrew Ransick, Nils O. Lindström, Jing Liu, Zhu Qin, Jin-Jin Guo, Gregory F. Alvarado, Albert D. Kim, Hannah G. Black, Junhyong Kim, Andrew P. McMahon

## Abstract

Chronic kidney disease affects 10% of the population with notable differences in ethnic and sex-related susceptibility to kidney injury and disease. Kidney dysfunction leads to significant morbidity and mortality, and chronic disease in other organ systems. A mouse organ-centered understanding underlies rapid progress in human disease modeling, and cellular approaches to repair damaged systems. To enhance an understanding of the mammalian kidney, we combined anatomy-guided single cell RNA sequencing of the adult male and female mouse kidney with in situ expression studies and cell lineage tracing. These studies reveal cell diversity and marked sex differences, distinct organization and cell composition of nephrons dependent on the time of nephron specification, and lineage convergence, in which contiguous functionally-related cell types are specified from nephron and collecting system progenitor populations. A searchable database integrating findings to highlight gene-cell relationships in a normal anatomical framework will facilitate study of the mammalian kidney.

## Introduction

The paired mammalian kidneys maintain homeostasis of the body’s fluids, remove metabolic waste products, control blood pressure and blood cell composition, and bone mineralization (Nielsen et al., 2012). Approximately 20% of cardiac output is directed to the kidney where a plasma filtrate passes through the glomerular vasculature into the proximal lumen of epithelial nephrons (Munger et al., 2012). A mouse kidney contains around 14,000 nephrons (Bonvalet et al., 1977), the human kidney around a million (Hughson et al., 2003). All nephrons develop from a mesenchymal, progenitor pool through reiterative inductive processes over days (mouse) or weeks (human) (McMahon, 2016). Each nephron comprises multiple segments with distinct physiological activities dependent on the repertoire of channels, transporters and enzymes within the segment (Nielsen et al., 2012).Nephrons align along a radial, cortical-medullary axis of functional symmetry, allowing for properties such as the concentration of urine. The nephron connects to the arborized ureteric epithelial network of the collecting system which has a distinct developmental origin to the nephron (McMahon, 2016). Specialized cells of the collecting duct balance systemic water, salt and pH levels (Pearce et al., 2015; Roy et al., 2015), with urine passing from the collecting duct through the ureter to the bladder for storage before excretion.

An estimated 5-10 million people die each year from kidney injury and disease (Luyckx et al., 2018). Though a kidney transplant is an effective solution, donor supply is well short of demand and transplant is not an option for many (Crews et al., 2019). End-stage renal disease is treated with dialysis, but dialysis is associated with high rates of morbidity and mortality (Mandel et al., 2016). Synthetic approaches to generate new kidney structures require a detailed understanding of kidney cell types and their actions (Oxburgh et al., 2017). Further, sex differences suggest male and female kidneys respond differently to injury and disease. For example, female mice and humans are more resistant to ischemia or ER-stress-invoked renal injury (Hodeify, 2013; Park et al., 2004; Aufhauser et al., 2016). Single nucleus and single cell RNA-sequencing (scRNA-seq) have provided new insight into the cellular make-up of mammalian organ systems including the kidney (Wu et al., 2019; Habib et al., 2017; Macosko et al., 2015; Adam et al., 2017; Chen et al., 2017; Park et al., 2018; Karaiskos et al., 2018; Han et al.,2018). However, kidney studies have been largely male-centered and lack insight into higher order organization of cell types critical to kidney function. Here, we provide new cellular insights into the organization, origins, diversity and diversity-generating processes for the male and female mouse kidney, providing access to these data through a searchable, anatomically-registered database.

## Results

### Anatomy directed profiling of the mouse kidney

Several considerations shaped our approach to single cell analysis of the kidney schematized in Figure 1a. First, scRNA-seq analysis should examine both sexes. Second, subdividing the kidney into distinct anatomically-related zones prior to cell dissociation would facilitate zonal optimization of representative cell recovery and cell mapping to the functional anatomy of the radially configured kidney cell organization. Third, the depth and quality of scRNA-seq data should permit identification of novel cell heterogeneity while minimizing tissue dissociation artefacts. Fourth, secondary validation of scRNA-seq-directed predictions would culminate in the creation of an anatomy-based, web-searchable view of these data.

**Figure 1.**
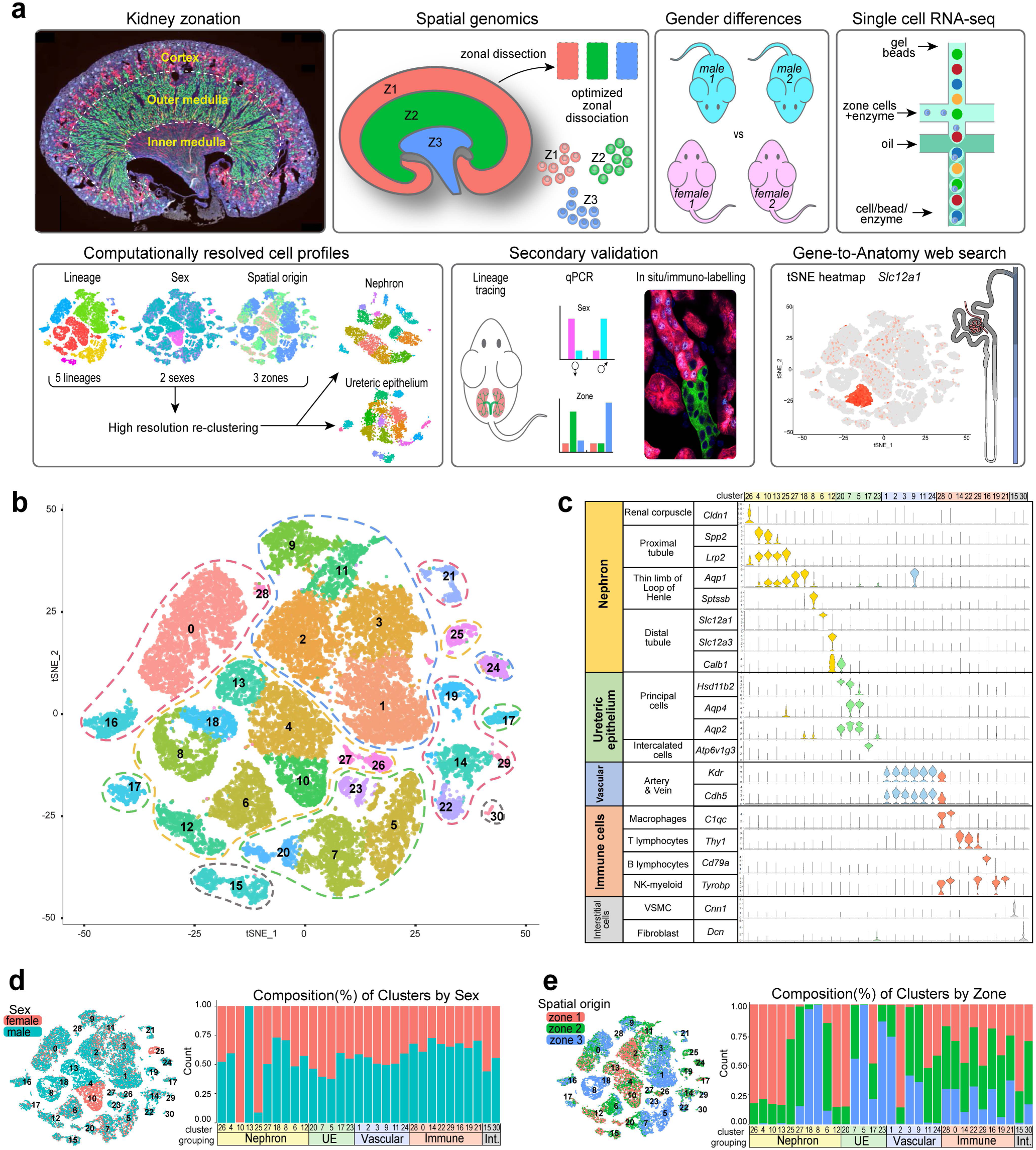
An anatomy-directed map of cell diversity in the adult mouse kidney. **a**, Overview of methodology, data collection and analyses. **b**, tSNE projection of adult mouse kidney scRNA-seq dataset annotated to five major cell-type groupings (dashed lines): nephron – yellow, ureteric – green, vascular – blue, immune – red, interstitial – gray. **c**, Violin plots of marker genes arranged by group and cell type, cluster numbers at top. **d-e**, tSNE and stacked barplots of adult mouse kidney dataset in (b), illustrating the distribution (left) and composition (right) of the clusters regarding sex (d) or zone (e). *See also Figure S1*

Single cell dissociation was optimized for micro-dissected zones of adult male and female mouse kidneys to obtain cell samples that conserved regional features of cell organization and cell diversity (Figure 1a). Varying the temperature (Figure S1a) and time (Figure S1b) of enzymatic digestion with a modified psychrophilic-protease protocol (Adam et al., 2017) enabled representative recovery of viable cells with minimal stress signatures. However, as noted by others (Park et al., 2018), podocytes, which show particularly strong cell interactions, were underrepresented. Single cell suspensions from the cortex (Zone 1), outer medulla (Zone 2) and inner medulla (Zone 3) of two adult (8-9 week) male and female C57BL6/J mice were processed through the 10X Genomics Chromium platform with Illumina Hi-Seq sequencing of barcoded single-cell libraries. The post-alignment sequencing metrics show an average of 717 million sequencing reads across 10,178 cells per kidney, with a median average of 1,395 genes/cell (Figure S1c).

Computational analysis for differentially expressed genes merged all 12 gene/barcode matrices, each annotated for replicate, zone and sex of origin. After filtering weak and outlying profiles (Figure S1d), profiles of 31,265 cells where analyzed in the Seurat R package by an unsupervised clustering: the data are displayed as a two-dimensional tSNE plot (Figure 1b) and tabulated lists of the most differentially expressed genes (Table S1). The 30 distinct cell profiles resulting from this primary analysis were annotated by matching enriched gene sets with kidney cell-type specific markers (Chen et al., 2017; Han et al., 2018; McMahon et al., 2008; Lee et al., 2015), grouping the dataset into five major lineage compartments (Figure 1c). Replicates were similarly represented in most clusters (Figure S1e), although a marked sex bias was evident in several proximal tubule-associated clusters (Figure 1d). Zonal analysis was in good agreement with expectations: proximal tubules, distal convoluted tubules and the cortical collecting duct were recovered from zone 1, while long thin loops of Henle and inner medullary collecting duct localized to zone 3 (Figure 1e). In summary, initial clustering analysis demonstrated reproducibility throughout the dataset while highlighting sex-related differences in the nephron and spatial diversity amongst the cell clusters.

Subsequent analyses focused on improving an understanding of nephron (N) and ureteric epithelium (UE) compartments given the key roles of this continuous epithelial network in renal physiology. Differentiation of N and UE progenitors in fetal and early postnatal development generates epithelial segments with distinct functions that are radially aligned along the cortical-medullary axis of the kidney (McMahon, 2016). The primary renal filtrate generated in the renal corpuscle passes through the proximal tubules, thin limbs of the loop of Henle and distal tubule segments to the UE-derived collecting system (Figure 2a). The origins of the connecting segment joining the nephron and ureteric networks has not been definitely reconciled. Developmental studies have suggested a UE origin (Howie et al., 1993), a N-progenitor origin (Kobayashi et al., 2008; Georgas et al., 2009), or a more complex hybrid with contributions from both lineages (Schmitt et al, 1999). Recognizing a difficulty in lineage assignment, clusters 12 (*Calb1^+^/Slc12a3^+^)* and 20 (*Calb1^+^/Hsd11b2^+^/Aqp2^+^)*, which both display high levels of the connecting segment marker *Calb1*, were annotated to N and UE groupings, respectively (Figure 1b-c).

**Figure 2.**
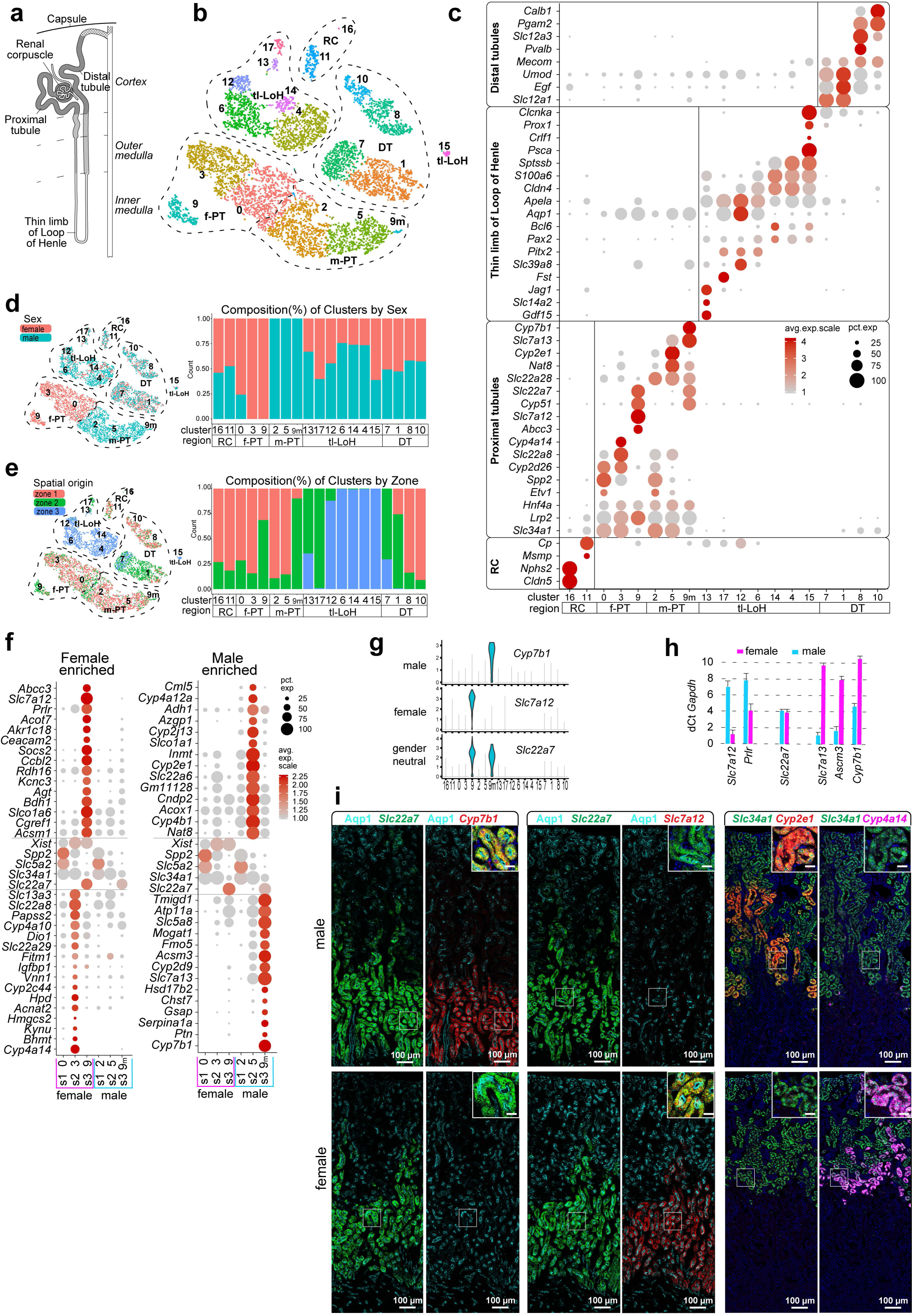
Sexual diversity within the nephron of the adult mouse kidney. **a**, Schematic showing a nephron (shaded) connecting (stippled) to the UE (un-shaded). **b**, tSNE projection of re-clustered N dataset using cells from primary annotation (yellow in Fig 1b). RC - renal corpuscle, tl-LoH - thin limbs-Loop of Henle, DT - distal tubules, PT - proximal tubules, male (m) or female (f). **c**, Dot plot of cluster enriched gene expression. **d-e**, tSNE and stacked barplots of N dataset illustrating the distribution (left) and composition (right) of clusters by sex (d) or zone (e). **f**, Dot plots of selected sex-biased gene expression in PT clusters. **g**, Violin plots of sex specific PT S3 genes. **h**, Quantitative PCR of sex-biased gene expression in the adult kidney. **i**, RNAscope in situs validate sex-biased gene expression in S3 and S2 segments of the proximal tubule. Inset scale bar = 20µm. *See also Figure S2*

### Sex-related diversity in nephron segments

Re-clustering of the assigned nephron cell types with Seurat through several iterations resolved 18 clusters (Figure 2b; Figure S2a). Cluster enriched gene expression was used to identify cell clusters (Figure 2c; Table S2). Strikingly, while most clusters were evenly represented by all kidney samples (Figure S2b), all proximal tubule clusters separated by sex (Figure 2d). Consistent with this finding, analysis of sex differences in whole kidney studies have highlighted male and female differences mapping to proximal tubule regions (Rinn et al., 2004). Clusters 0, 2, 3, 5, primarily located in the cortex (Figure 2e.), separated directly on the basis of sex, cluster 9 formed large female (298 cells) and small male (32 cells) groupings, secondarily designated as 9 and 9m, respectively (Figure 2b-d; Figure S2a-b). A sex bias in gene expression was observed across many gene categories linked to proximal tubule functions including organic anion (*Slc22a6, Slc22a8*) and amino acid (*Slc7a12, Slc7a13*) transport, and drug (*Fmo5*, *Cyp4a14),* cholesterol *(Cyp7b1*) and hormone (*Agt, Hsd17b2*) metabolism (Curthoys and Moe, 2014)

To map sexually dimorphic domains, we identified genes with enriched expression in male and female (*Xist^+^*) clusters, comparing their expression with genes known to highlight the segmental organization of the proximal tubule that exhibited comparable expression between the sexes: *Spp2/Slc5a2* (S1), *Slc34a1* (S1-2) and *Slc22a7* (S3) (Figure 2f). The highest disparity in gene activity between the sexes was in the S3 region. As examples, *Slc7a12* and *Cyp7b1* transcripts localized exclusively to female (cluster 9) and male (cluster 9m) subsets, respectively, of *Slc22a7*^+^ S3 cells (Figure 2g). Quantitative PCR analysis of cDNA from unfractionated adult mouse kidneys corroborated sexually dimorphic gene activity (Figure 2h). To visualize gene expression directly in PT segments, we combined uniquely labelled RNAscope probes and performed *in situ* hybridization to adult male and female kidney sections. As predicted, *Slc7a12* showed female-restricted expression and *Cyp7b1* male-restricted expression in the *Slc22a7*^+^ S3 subset of the Aqp1^+^ proximal tubule (Figure 2i), while male-restricted expression of *Acsm3* and female-restricted expression of *Prlr* extends at lower levels into the S2 domain (Figure S2c). Further, co-analysis of *Slc34a1*, *Cyp2e1* and *Cyp4a14* transcripts highlighted sex-restricted activity of *Cyp2e1* (male only) and *Cyp4a14* (female only) in the S2 subset of *Slc34a1*+ PT regions (Figure 2i). Interestingly, S2 and S3 sex-specific expression domains form a tight, non-overlapping boundary at the S2-S3 junction (Figure S2d). Gene Ontology analysis comparing sex differences in the entire proximal tubule and S3 regions specifically highlighted differences in small molecule, lipid and organic acid metabolism (Figure S2e; Table S3).

These findings are in good agreement with earlier insights identifying sex-related differences in the kidney predominantly through whole kidney studies (Rinn et al., 2004; Si et al., 2009; Sabolić et al 2007) but significantly extends the repertoire of genes with sex specific activity as well as resolving expressing cell types. At present, there is no comparable scRNA-seq data for the human kidney though kidney biopsy analysis points to sex differences (Si et al., 2009). However, mice and humans both show a male-biased susceptibility to ischemia invoked kidney injury where the S3 region is particularly vulnerable to injury (Park et al., 2004; Neugarten et al., 2018) A susceptibility to stress induced cell death may explain why male S3 cells were markedly underrepresented in the scRNA-seq profile (∼10% of expected; Figure S2a, Fig 2i).

Additional studies will be required to determine the significance of sex differences and a human relevance, some differences are particularly intriguing. For example, the prolactin hormone receptor (*Prlr*) is preferentially expressed in the female S3 region (Figure 2f, h; Figure S2c). These data suggest a Prlr-driven regulation of renal function to the benefit of mother, offspring, or both. In the mammal, prolactin produced by the pregnant and nursing female regulates milk production in the mammary gland and hypothalamic-directed behaviors through Prlr. Functional studies in the human and rat have linked prolactin to salt and water control in the proximal tubule and evolutionary studies have suggested a link between prolactin and sodium regulation in the salt to fresh water transition of fish (Ibarra et al., 2005; Freeman et al., 2000). Additional female S3 enriched genes are linked to hormone metabolism (including *Dio1*, *Ttr*, *Akr1c18*, *Gm4450*, *Spp1* and *Rdh16*) (Piekorz et al., 2005) and kidney damage or disease (including *Spp1, Lrp2* and *Cubn*) (Lorenzen et al., 2008; Pei et al., 2016).

### Spatial and temporal diversity of nephron cells

The 14,000 nephrons of the mouse kidney are generated from a progenitor population over a protracted period of fetal and neonatal development through a reiterative inductive process (McMahon, 2016). Juxtamedullary nephrons have renal corpuscles close to the cortical-medullary boundary and project long loops of Henle deep into the inner medullary domain while cortical nephrons have peripheral cortically localized renal corpuscles and project short loops of Henle restricted to the outer medulla (Figure 3a). The long and short loops have distinct physiological actions, with urine concentration associated with the deep medullary loops of the juxtamedullary nephrons (Dantzler et al., 2011; Jamison, 1987). A limited understanding of the diversity of nephron types, their origin, and the cell diversity within the loops of Henle, makes annotating this region a particular challenge.

**Figure 3.**
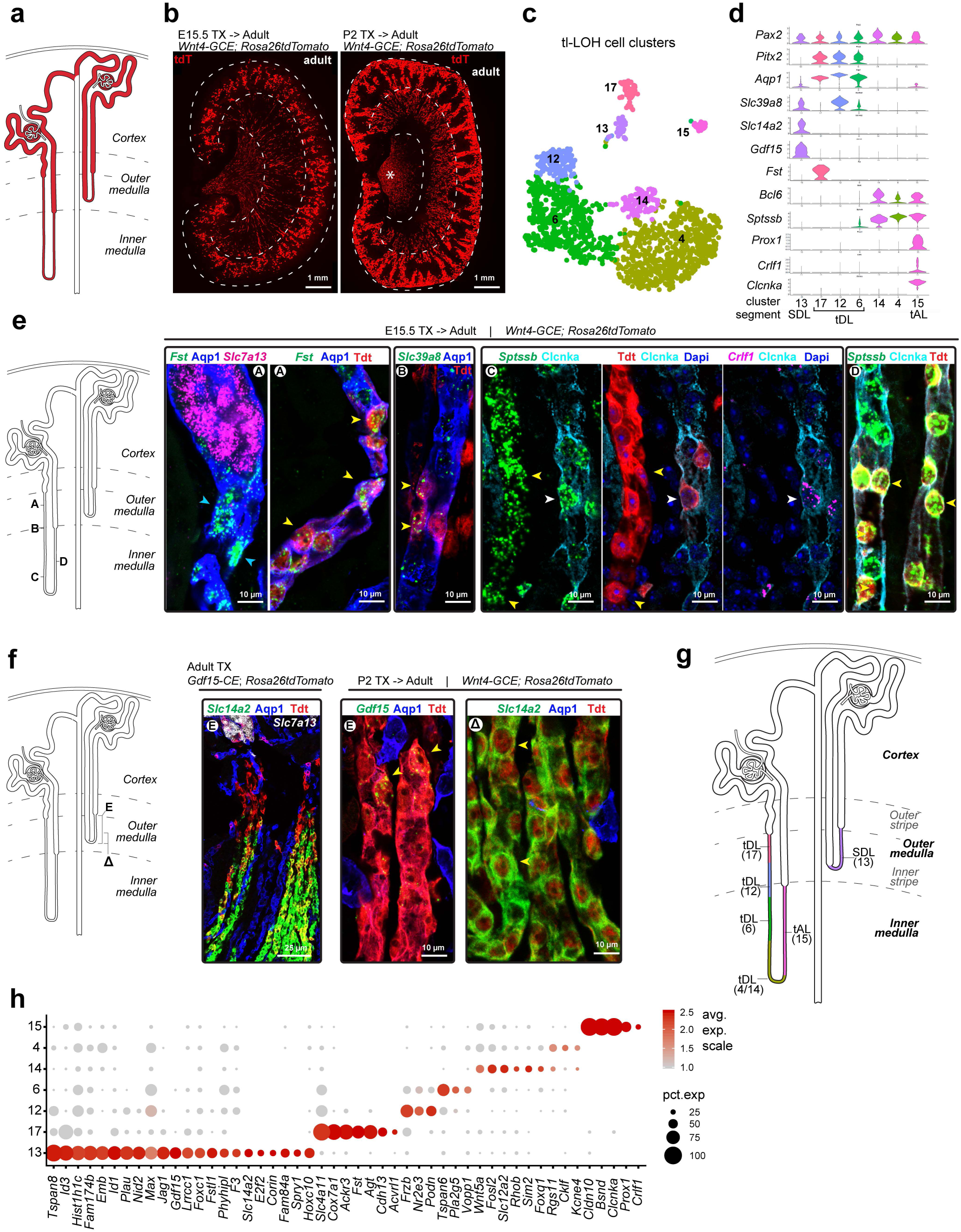
Divergent cell fates and anatomies in the thin limbs of loop of Henle linked to developmental timing. **a**, Schematic of juxtamedullary (left) and cortical nephrons (right); dashes demarcate boundaries of cortico-medullary zones. **b**, Views of adult kidneys showing nephron populations formed at ∼E15.5 (left) or ∼P2 (right) labelled using inducible lineage-tracing; see Methods for details; asterisk in (b) - non-nephron interstitium. **c-d**, Thin limbs of Loop of Henle (tl-LoH) cell clusters and violin plots of segment-identifying genes; cluster 15 repositioned here - compare with Fig 2b. SDL - short descending limb; tDL - thin descending limb; tAL - thin ascending limb; **e-f**, RNAscope validation of segmental markers for tl-LoH in juxtamedullary and cortical nephrons (e-f, respectively); Arrowheads: A, blue: *Fst*^+^/Apq1^+^; yellow: *Fst*^+^/Apq1^+^/tdT^+^; B, *Slc39a8*^+^/Apq1^+^/tdT^+^; C, yellow: *Sptssb*^+^/*Crlf1*^-^/Clcnka^-^/tdT; white: *Sptssb*^+^/*Crlf1*^+^/Clcnka^+^/tdT^+^; D, *Sptssb*^+^/Clcnka^+^/tdT^+^; E, *Gdf15*^+^/Aqp1^-^/tdT^+^; Δ, *Slc14a2*^+^/Aqp1^-^/tdT^+^. **g**, Schematic proposing anatomical organization and cell type identity of clusters shown in (c). **h**. Dot plot showing selected genes enriched in individual tl-LoH clusters. *See also Figure S3*

To address the question of nephron origins, a *Wnt4^CRE-ERTM^* mouse strain (Kobayashi et al, 2008) was used to activate a tandem tomato (tdT) fluorescent lineage tracer in nephron precursors at embryonic day 15.5, or postnatal day 2 (Figure 3b). Early labelling marked juxtamedullary nephrons and late marking, cortical nephrons, consistent with a spatial temporal progression in assembly of the kidney cortex (Figure 3b). Next, we examined clusters of profiled cells tentatively assigned to the thin limb of the loops of Henle (Figure 3c), identifying genes with expression enriched in each cluster (Figure 3d), then mapped gene activity to tdT^+^ cells in early- and late-formed nephron types (Figure 3e-f).

Juxtamedullary nephrons transitioned in the outer medulla from Aqp1^+^/*Slc7a13*^+^ cells in the S3 region in the outer stripe to Aqp1^+^/*Fst*^+^ cells in thin descending limb of the loop of Henle in the inner stripe (cluster17, Figure 3c-e; Figure S3a). From the outer to inner medulla, the descending limb transitions sequentially through several distinct cell types: Aqp1^+^/*Slc39a8*^high^ (cluster 12), Aqp1^+^/*Slc39a8*^low^ (cluster 6) and Aqp1^-^/*Sptssb*^+^ (clusters 4 and 14) (Figure 3c-e; Figure S3a). Clusters 4 and 14 at the transition from descending to ascending limb of the loop of Henle were not resolved further though *Sptssb* marked both Clcnka- and Clcnka+ cells within tdT+ thin limbs (Figure S3a). Clcnka^+^/*Sptssb*^+^/*Crlf1*^+^ (cluster 15) cells generated the ascending thin limb up to its junction with distal tubule segments at the outer medullary border (Figure 3c-e; Figure S3a).

In cortical nephrons, Aqp1^+^/*Slc7a13*^+^ S3 cells in the outer stripe of the outer medulla transitioned to Aqp1^-^/*Gdf15+* cells (cluster 13) identified through cell fate mapping with a *Gdf15^CRE-ERTM^* mouse strain (unpublished APM; Figure 3f; Figure S3b). The Gdf15^+^ region transitions to a *Slc14a2^+^* region within the inner stripe of the outer medulla before connecting to the thick ascending limb of the loop of Henle (Figure 3c-d, f; Figure S3b). As expected, no overlap was observed between juxtamedullary (*Fst^+^*) and cortical (Gdf15-tdT^+^) nephrons (Figure S3b).

A schematic model highlights the relationship of cell diversity in the loop of Henle to nephron type (Figure 3g). Acquisition of loop of Henle functions were critical to the successful radiation of mammalian species (Jamison, 1987). Low nephron count is linked to kidney disease (Hughson et al., 2003) and reduced nephron formation is predicted to preferentially impact formation of later arising cortical nephrons. Gene enrichment analysis gives insights into distinct activities and potential regulatory actions associated with specific cell types of juxtamedullary and cortical nephrons (Figure 3h). Focusing on the thin limb of the loop of Henle region of late forming cortical nephrons (cluster 13), analysis shows enriched expression of genes encoding transcriptional regulators (Id1, Id3, Foxc1 and E2f2) and signaling components (Jag1, Gdf15, Spry1 and Fstl1), suggesting mechanisms for the control of regional cell identity and local cell interactions (Figure 3h). Examining disease linkage, *Corin* expression, which is quite specific to the cortical nephron-specific cell population, encodes a serine peptidase critical to the regulation of blood volume and blood pressure. Human variants in Corin are linked to hypertension, cardiac hypertrophy and pre-eclampsia (Li et al., 2017).

### Spatial diversity in the collecting system

The ureteric epithelium (UE) of the collecting system (Figure 4a) has a temporally and spatially distinct origin from the nephron progenitor forming lineage (Taguchi et al., 2014). A re-clustering with Seurat of the annotated UE cell profiles, including the cortical connecting tubule from the primary dataset (Figure 1b,c), resolved 16 clusters ranging from 61 to 543 cells (Figure 4b; Figure S4a). Gene enrichment analysis (Figure 4c; Table S4) coupled with a zonal analysis of the expected distribution of key cell types (Figure 4d) facilitated identification of each cell cluster.

**Figure 4.**
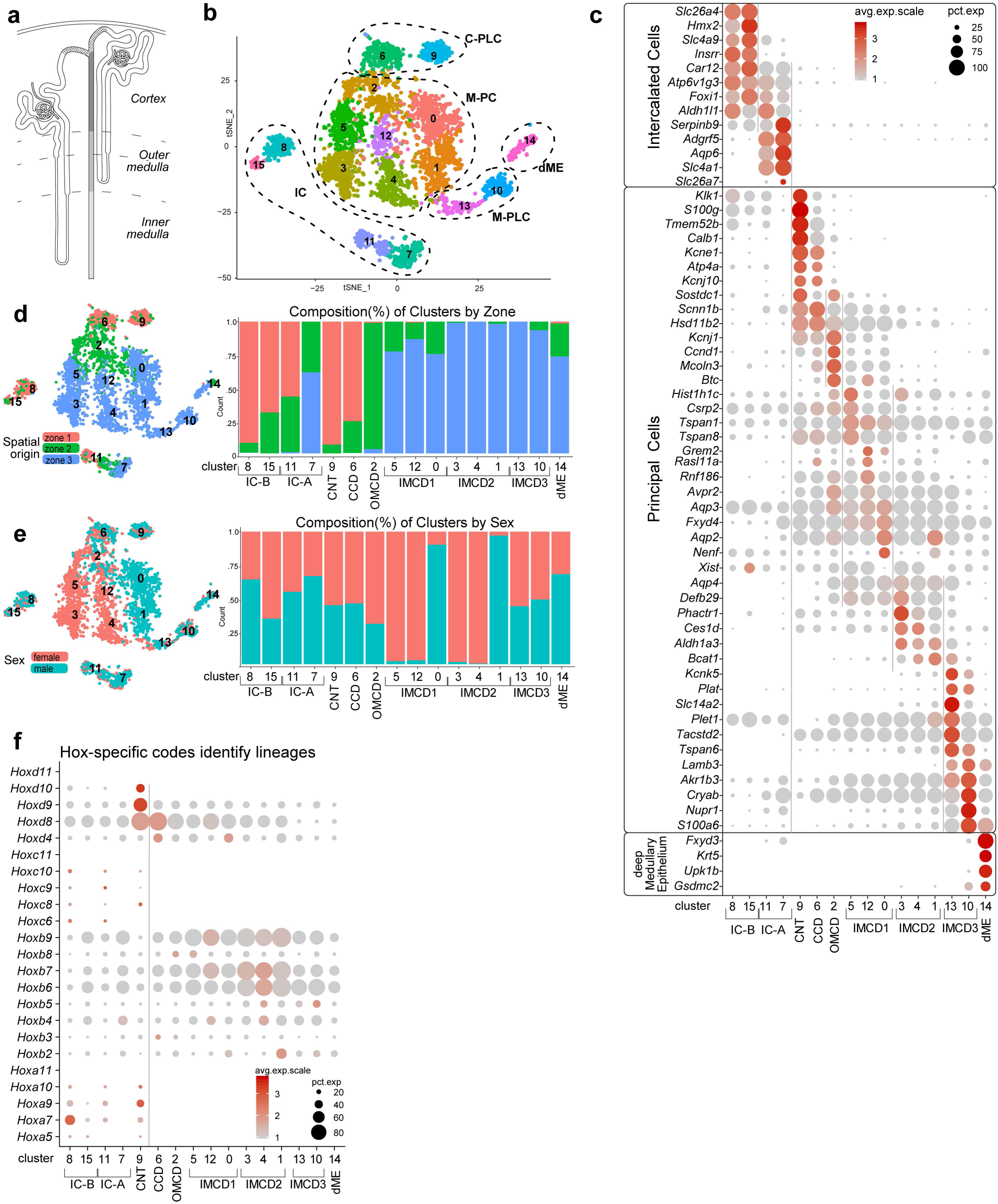
Anatomical, sex and cell diversity of ureteric epithelial groupings. **a**, Schematic showing nephron (unshaded) to UE (shaded) transition through connecting segment (stippled). **b**, tSNE projection of re-clustered UE dataset using cells from primary annotation (green in Figure 1b); C- and M-PLC, cortical and medullary principal-like cells; M-PC, medullary principal cells. **c**, Dot plots of cluster-enriched gene expression, annotating clusters as intercalated cells (IC) type A and B, connecting tubule (CNT), cortical collecting duct (CCD), outer and inner medullary collecting duct (OMCD, IMCD types1-3), and deep medullary epithelium (dME). **d-e**, tSNEs and barplots illustrating the distribution (left) and composition (right) of the cell clusters with respect to zonation (d) and sex (e). **f**, Dot plot summarizing *Hox*-gene expression across the UE dataset cell clusters. *See also Figure S4*

A single Aqp2^+^ and Aqp3^+^/Aqp4^+^ principal cell type has been thought to regulate water and salt levels in response to hormonal input. However, a striking diversity was observed amongst cells annotated with PC features (Figure 4b-c). The diversity relates at least in part to position along the cortical-medullary axis (Figure 4d). Analysis of specific gene profiles amongst these clusters (Figure 4c) demonstrates cortical clusters 9 and 6 and medullary clusters 13 and 10 share PC-like signatures with medullary clusters 2, 5, 12, 0, 3, 4, 1 but showed markedly distinct profiles from these PCs. We designate clusters 9, 6, 13 and 10 as principal-like cells (PLC) where *Kcne1* and *Atp4a* distinguishes cortical PLC clusters, and *Lamb3* and *Plat* PLC clusters of the inner medullary collecting duct (Figure 4b-c; Table S4). In addition to a positional bias, a marked sex bias was also observed amongst the PC clusters (Figure 4e). Further, whereas the two male samples replicate well, the two female samples show a markedly more divergent clustering (Figure S4b). Collectively, the data suggest marked regional and sex-related differences in the transcriptional state of PCs in the mouse kidney. Interestingly, the greater diversity observed between female replicates is not likely to be a technical artefact as no bias was observed in other clusters. Variability in the stage of estrus in the adult females undergoing reproductive cycling could underlie female tissue variability.

Intercalated cells play an important role in acid-base and electrolyte homeostasis (Roy et al., 2015). Three intercalated cell types are widely recognized: Slc4a1^+^ IC-A cells which distribute along the entire cortico-medullary axis, Slc26a4^+^ IC-B cells localized exclusively to cortical regions (Chen et al., 2017) and a distinct cortical “nonA-nonB” IC cell-type distinguished on the basis of cell shape and the distinct membrane polarity of transmembrane proteins (Weiner and Verlander, 2011, 2017). Non-supervised clustering identified two distinct IC-A (clusters 7 and 11) and two IC-B (clusters 8 and 15) populations exhibiting expected zonal restrictions (Figure 4b, d). Examining *Hox* gene expression amongst IC cell clusters provided additional insight into these clusters. *Hox* gene activity gives a positional history of cell origins, with higher numbered *Hox* genes expressed by cells in increasingly more posterior positions along the body axis (Mallo, 2018). Deregulated *Hox* gene activity results in dramatic kidney phenotypes (Wellik et al., 2002; Drake et al., 2018). Only cortical IC-A cluster 11, IC-B cluster 8 and PLC cluster 9 displayed a *Hox10* paralog expression signature (Figure 4f) consistent with IC and PLC types arising independently from nephron (*Hox10^+^*) and ureteric (*Hox10^-^*) lineages, reflecting the temporal-spatial sequence in the establishment of these lineages in embryonic development (Taguchi and Nishinakamura, 2017).

### Lineage diversity at the nephron-collecting system junction

To address the origins of IC and PLC/PC lineages directly, we visualized nephron and collecting system junctions by genetically labelling all cells of nephron progenitor origin with tdT (Figure 5a; Six2^GFP-CRE^; *Rosa26^tdTomato^* strain) (Kobayashi et al., 2008; Madisen et al., 2010) and all ureteric derivatives with Venus protein (Figure 5a; tgHoxb7-Venus strain) (Chi et al., 2009). Junctions displayed different patterns of tdT^+^ and Venus^+^ cell-intermingling but with a clear separation of each marker protein (Figure 5b-d; Figure S5a-d). Infrequent tdT^+^/Venus^+^ co-labelled cells close to the border zone may reflect rare instances of cell fusion (Figure 5c). As predicted from the single cell data, *Hoxd10*^+^ cells were always tdT^+^ indicating these cells arise from the nephron lineage (Figure 5d). Immuno-identification of Aqp2^+^ PLCs and Atp6v1b1^+^ ICs showed tdT^+^ and Venus+ labelling of both cell classes (Figure 5e; Figure S5a). Thus, nephron and ureteric lineages contribute to both IC and PLC populations in the junction region.

**Figure 5.**
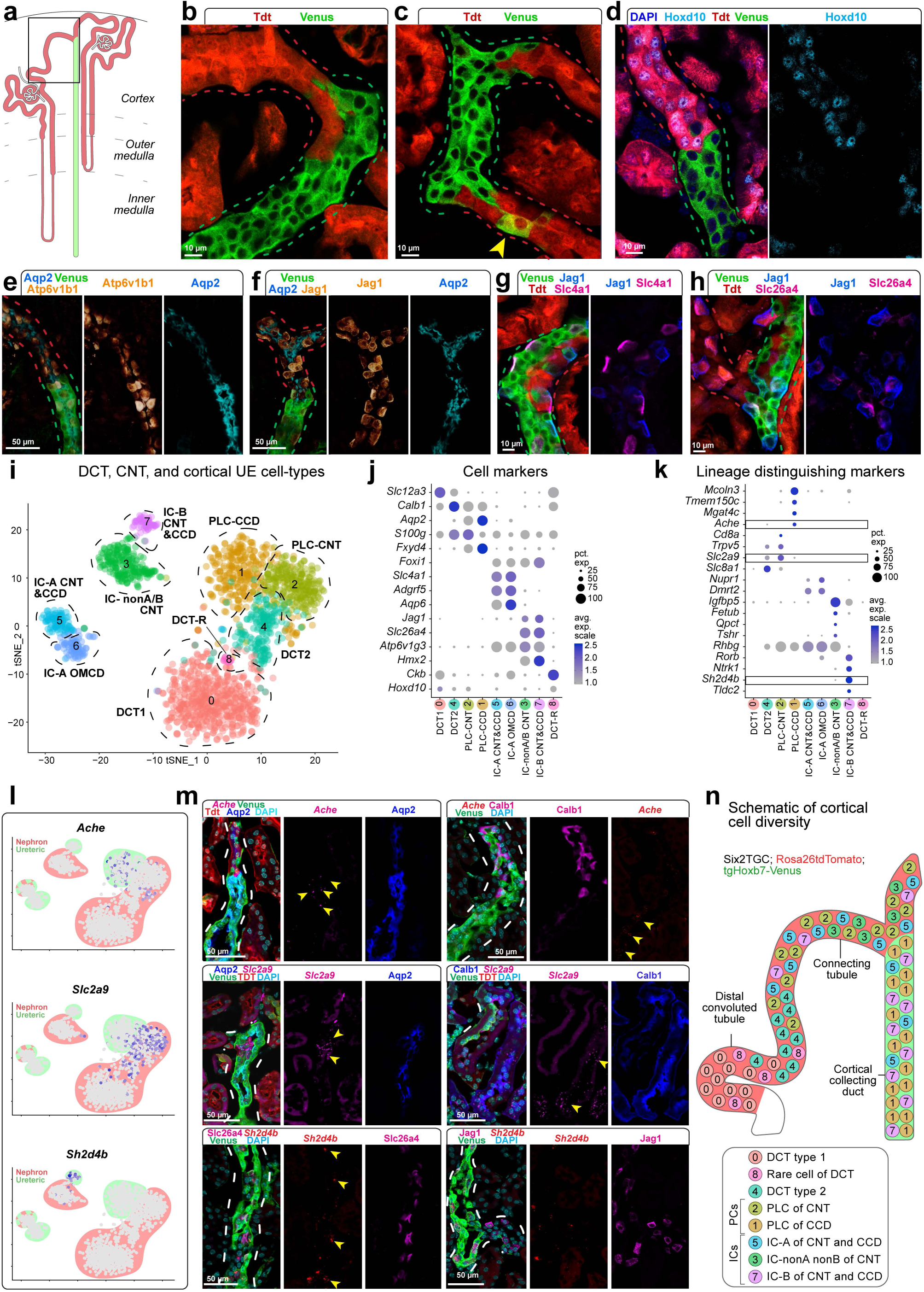
Dual origin for cortical principal and intercalated cell types. **a**, Schematic showing nephron (red) and ureteric (green) lineage labeling and region of focus (boxed). **b-c**, Lineage-tracing identifying nephron connections with the UE; arrowhead, co-labelled cell. **d-h**, Distribution of positional (Hoxd10), principal (Aqp2) and intercalated (Atp6v1b1, Jag1, Slc4a1 and Slc26a4) cell markers by immunostaining. **i**, tSNE projection of re-clustered cortical epithelial cell types from N and UE datasets. **j**, Dot plot of cluster enriched gene expression. **k**, Dot plot of lineage enriched gene sets; boxes show genes validated by RNAscope *in situ*. **l-m**, Feature plots and RNAscope validation of lineage distinguishing cell markers; yellow arrowheads, transcripts; dashed lines, tubule outlines. **n**, Schematic summary of cell types of the nephron and UE connection. *See also Figure S5*

Previous reports have demonstrated PC and IC fates are dependent on Jag1 regulation of the Notch pathway and the transcriptional actions of Tfcp2l1 (Werth et al., 2017; Jeong et al., 2009; Bielesz et al., 2010). Analysis of Jag1 (IC-B restricted) and Tfcp2l1 (broad ICs and weak PLC) showed the expected activity independent of their origin, suggesting similar mechanisms determining PLC and IC fates from ureteric and nephron lineages (Figure 5f; Figure S5b-c). Similarly, proteins specifically demarcating IC-A or IC-B cell types (Slc4a1 and Slc26a4/Jag1, respectively) were present in cells originating from both lineages (Figure 5g-h).

To facilitate resolving cell types at the junctions, all cortical cells from the distal nephron and ureteric lineages were sub-clustered (Figure 5i) and differentially expressed gene markers identified for each cluster (Figure 5j; Figure S5e-i; Table S5). In addition to providing further insight into IC cell populations, sub-clustering identified a novel subset of Ckb^+^ cells within the Slc12a3^+^ distal convoluted tubule of the nephron (Figure S5j). PLCs separated into clusters that were predominantly of nephron (*Aqp2^+^*/*Hoxd10*^+^ cells, cluster 2) or ureteric (*Aqp2^+^*/*Hoxd10*^-^ cells, cluster 1) origin (Figure 5i-j; Figure S5h-i). Cortical IC-A cell types formed a single cluster (#5) of *Slc4a1*^+^/*Hoxd10*^+^ IC-A types of nephron origin and likely UE-derived *Slc4a1*^+^/*Hoxd10*^-^ (Figure 5i-j; Figure S5h-i); a distinct grouping from medullary IC-A types (cluster 6; Figure 5i-j). IC-B cells were distinguished by expression of the anion transporter Slc26a4 (Park et al., 2018). *Slc26a4*^+^ cells segregated into nephron-derived (*Hoxd10^+^*) and ureteric-derived clusters 3 and 7, respectively (Figure 5i-j; Figure S5h-i). Interestingly, expression of the ammonium transporter *Rhbg* distinguishes clusters 3 and 7. Co-expression of *Slc26a4* and *Rhbg* is a feature of nonA-nonB intercalated cells (Weiner and Verlander, 2011). Direct immunolocalization demonstrated Slc26a4^+^/Rhbg^+^ nonA-nonB ICs were exclusively derived from the nephrogenic lineage (Figure S5k). In addition, approximately 20% of Slc26a4^+^/tdT^+^ cells were negative for Rhbg, indicating nephron progenitors also generate IC-B cells.

To find unique gene identifiers for cell types within the junction region, we performed correlation gene expression analyses for *Aqp2* and *Slc26a4* comparing *Aqp2*^+^/*Hoxd10*^+^ with *Aqp2*^+^/*Hoxd10*^-^ cells and *Slc26a4*^+^/*Hoxd10*^+^ with *Slc26a4*^+^/*Hoxd10*^-^ cells, followed by a secondary correlation amongst selected genes (Figure 5k-l). RNAscope *in situ* hybridization of predicted PLC markers validated nephron lineage enrichment for *Slc2a9* expression and ureteric lineage enrichment for *Ache* expression, while *Sh2db4* distinguished IC-B cells from other cell types (Figure 5m; Figure S5l-m).

In summary, these data point to a diversity of cell types and cell origins where nephrons connect to the collecting system, a region critical for hormone-dependent regulation of sodium and calcium resorption (Markadieu et al., 2011)(Figure 5n). Nephron progenitors generate PLCs, IC-A, IC-B and nonA-nonB IC cells and cortical ureteric epithelial progenitors give rise to PLCs, IC-A and IC-B cells. Cortical PLCs and IC-A from nephron and ureteric lineages are more similar to each other than to ureteric epithelial progenitor derived IC-A and PC/PLCs of the medullary region. The dual origins for PLCs and IC cell types is particularly intriguing. As mouse and human epithelial nephron precursors transition from a renal vesicle to an S-shape body physically connected to the ureteric epithelium, the distal region progressively adopts a transcriptional regulatory signature similar to the adjacent ureteric epithelium, including the activation of regulatory factors such as Tfcp2l1 that are essential for the specification of IC cell fates (Figure S6a-d) (Lindström et al., 2018a,b). Collectively, these data suggest an *in vivo* reprogramming of distal nephron identities to resemble those of the ureteric lineage. These findings have important implications for regenerative strategies to restore human kidney function (Oxburgh et al., 2017). The plasticity and natural cell heterogeneity in the distal nephron may obviate the need for co-development of ureteric epithelial-derived cell types.

## Discussion

Combining single cell analyses with genetic fate mapping generated novel insights into cell, regional, lineage and sex-related diversity in the mammalian kidney. Representative sampling of cell types and re-association of single cell data to the functional anatomy of any organ system is not trivial and the kidney’s complexity makes this a particular challenge. The microdissection and cell dissociation strategy adopted here enabled the optimization of regional cell recovery while conserving key features of the structural organization critical to anatomical mapping of cell diversity. There are many additional ways in which these data may be productively mined. As an example, *de novo* specification of kidney cell types will be facilitated by insight into cell-type enriched transcriptional determinants specifically enriched within nephron and UE cell clusters (Figure S7). An overview highlights the transcriptional relationship amongst cell clusters comprising proximal, medial and distal segments of the nephron, that likely reflect developmental relationships in the foundation of nephron segmentation (Figure S7a). Similar relationships are observed along the radial axis of the collecting epithelium (Figure S7b). While the focus here has been on cells directly interacting with the plasma filtrate, preliminary analysis of vascular and immune cell datasets suggests interesting new insights.

Current informatics platforms for scRNA-seq data analysis require some coding expertise. Further, the level of annotation of scRNA-seq data is often quite minimal, diverging from accepted ontologies, hindering convergent analysis across datasets. To ensure community access to these data, we generated a simple web-searchable database - Kidney Cell Explorer - to visualize gene expression patterns through tSNE analysis (Figure 6a-c) and appropriately annotated combined nephron and UE models (Figure 6d-f; https://cello.shinyapps.io/kidneycellexplorer/). These models combine individual cell cluster data to create a robust “metacell” for each anatomical ontology group (Figure 6f) comprising 9,000 to 15,000 detected genes (Figure 6g). A gene search enables individual gene expression data to be viewed within each primary and secondary cluster (Figure 6a-c) and as a heat map across juxtamedullary and cortical nephron models. Further, a batch search option allows the comparison of multiple genes across each “metacell” grouping (Figure 6e; Figure S8). As illustrative examples of the utility of this approach, we show the distribution of selected genes associated with hormonal regulation (Figure S8a), kidney disease (nephrotic syndrome; Figure S8b) and mediating the kidneys physiological activities (Figure S8c). Interestingly, high levels of *Pth1r* expression in podocytes points to a potential for off target effects of its bone stimulating ligand PTH in treating osteoporosis (Figure S8a). With respect to sex differences, surprisingly, female proximal tubule segments show higher levels of expression of testosterone receptor (Ar) and growth hormone receptor (*Ghr*) but lower levels for estrogen receptor (*Esr1*) than male segments. Further, expression of transcripts encoding the key blood pressure regulating enzymes *Ace* and *Ace2* show marked disparity between the sexes in proximal tubule regions (Figure S8a). Sex differences are also evident in the expression of multiple genes regulating uptake from, or secretion into, the plasma filtrate as it passes through the proximal tubule (Figure S8c). Examining cell origins of kidney disease, nephrotic syndrome genes highlight the podocyte as key target in agreement with recent studies (Park et al., 2018), but provide additional insight into disease genes predicted to impact the actions of many other cell types (Figure S8b).

**Figure 6.**
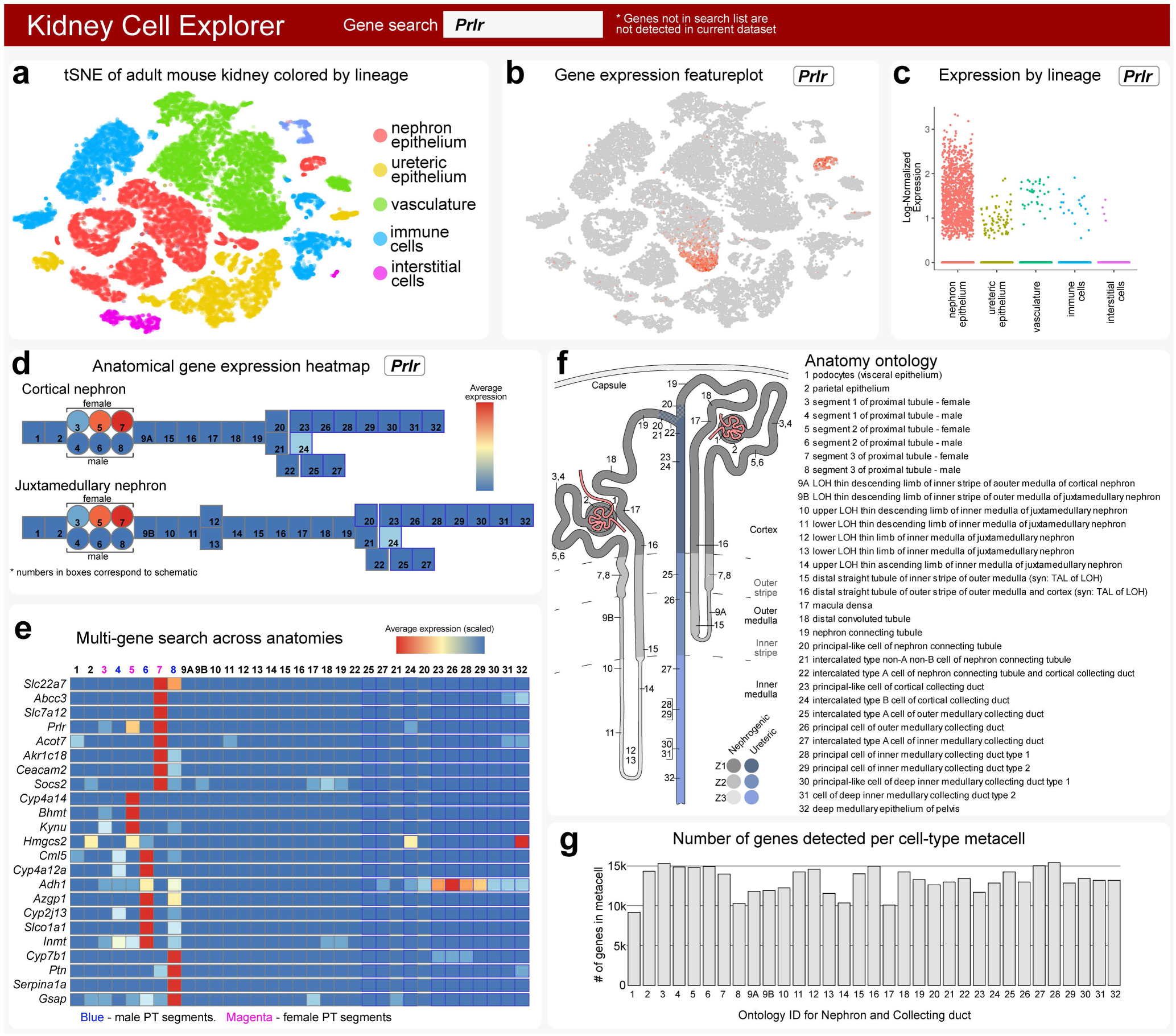
Kidney Cell Explorer views - a searchable map of cell diversity in the adult mouse kidney. **a**, Field enabling selection of tSNE plot for whole kidney, nephron or ureteric epithelium dataset views. **b**, Field producing a feature plot of selected gene in active scRNASeq dataset. **c**, Field producing scatterplot of selected gene as expressed across sex, spatial origin or lineage in whole kidney dataset. **d**, Heatmap of gene expression for a single selected gene in metacells arranged as cortical and juxtamedullary nephron models. **e**, Heat map of gene expression for multi-gene searches in metacells of nephron and collecting duct. **f**, Schematic map indicating anatomic position and ontology terms for metacells in (d). **g**, Number of genes detected in each metacell in (d). *See also Figure S8*

In conclusion, Kidney Cell Explorer will facilitate access to our data and analysis of these data by the academic and clinical research communities. Further, the overall strategy adopted here provides a working framework for the generation of a comprehensive cell atlas for the human kidney.

## Acknowledgments

The authors thank members of the McMahon and Smith laboratory for helpful comments and useful discussion on single cell analyses. Work in APM’s laboratory was partially supported by grants DK110792, DK054364 and an RBK partnership grant DK107350-01. JK and QZ were supported in part by RBK partnership grant DK107350-02S1. The Flow Cytometry Core is supported by NCI CCSG award number P30CA014089.

## Author contributions

AR, APM, NOL, and JL wrote the manuscript. AR, NOL, and JL assembled figures. AR, JL, NOL, HGB and APM designed and/or analyzed experiments. AR and ADK performed single-cell sequencing. JG, NOL, JL and GA performed secondary verification and follow up studies of single cell data predictions. QZ and JK built the interactive web site.

## Data deposition

The single cell RNA sequencing data is deposited at GEO accession number: GEO: GSE129798.

## Methods

### Adult mouse kidney processing

Adult C57BL/6J mice (9-10-week; 2 males, 2 females) were euthanized by CO2 inhalation for 3.5 minutes, followed by transcardial perfusion with 10-15 ml cold Dulbecco’s phosphate-buffered saline (DPBS) to remove blood cells. Kidneys were harvested, the kidney capsules removed and ureter trimmed prior to dissection with a scalpel into tissue blocks specific for cortex (zone 1, Z1), outer medulla (zone 2, Z2) or inner medulla (zone 3, Z3). All steps were carried out in a petri dish on ice. A superficial slice (∼25% depth) was made to reveal the Z2-Z3 boundary, then a central segment containing all of Z3 was isolated by making two complete cuts diagonal to the corticomedullary axis (CMA) and through the Z2-Z3 boundary. The two outer pieces of kidney resulting from step 2 were placed cut-surface down and cut along the cortical-medullary axis into several thick slices. Each slice was cut into Z1 and Z2 portions. All tissue blocks were examined for zonal composition and trimmed where necessary. To obtain ∼50 mg of pure Z1 or Z2 tissue, several pieces were combined from one kidney. The relative compactness of the Z3 region led to pooling tissue from both kidneys for a yield of ∼25-30 mg tissue for M1_Z3, M2_Z3 and F2_Z3 samples; the F1_Z3 sample was ∼15 mg tissue from a single kidney. All tissue blocks were kept in dishes on ice until processed further.

### Adult mouse kidney cell dissociation

Dissociation of the adult mouse kidney into a representative set of single cells presented a significant challenge beyond protocols extracting cortical nephron progenitor cells from the surface of fetal and newborn mouse kidney (Lindström et al., 2018a). To avoid stress artefacts, we followed a ‘cold dissociation’ strategy, built around *Bacillus licheniformis* cold active protease (CAP), which had been successfully used to dissociate newborn (p1) mouse kidneys (Adam et al., 2017), monitoring cell yields by fluorescence-activated cell sorting (see below) and a quantitative PCR assessment of cDNA for cell-type specific markers comparing to undissociated tissue. Cell dissociation times with CAP (10 mg/ml) and collagenase, type 2 (2.5 mg/ml) were markedly improved from ice-cold dissociation (>1.5 hrs.) by incubating samples at 12-16^°^C (30-45 minutes) without introducing a significant transcriptional stress response signature (Figure S1a,b).

Adopting 12^°^C incubation in collagenase-supplemented CAP produced similar yields (within 2-fold range) for most canonical cell type markers with 30 (Z1), 45 (Z2), or 60 (Z3) minutes of digestion (Figure S1b). The final concentration of reagents in the DPBS was 2.5 mg/ml collagenase, type 2 [Worthington, #LS00417]; 7.5 mg/ml *B. licheniformis* protease [Creative Enzymes, NATE-0633] and 125 U/ml DNase I [Worthington, #LS002058]. At five minute intervals, each digestion was mixed twenty times with a wide bore pipette and tissue dissociation was monitored under a dissecting microscope. Digestions were terminated by adding an equal volume of 20% fetal bovine serum (FBS) in DPBS and transferring the tubes to ice. Cell suspensions were strained through a pre-wetted “coarse 40 µm” strainer (Falcon), washing the filter with an additional 1ml DPBS. The flow-through was pelleted at 1250 rpm in a swinging bucket centrifuge set to 6^°^C then resuspended in 3 ml cold AutoMACs Running Buffer (AMB, Miltenyi Biotec). After gentle resuspension, the cells were passed through a pre-wetted “fine mesh 40 µm” strainer (VWR). The resulting flow-through was pelleted as before, then resuspended in 0.35 ml of freshly prepared AMB containing nuclear dyes 14 uM DAPI and 5 uM DR then sorted on an ARIA II FACS at a low flow rate, using gates for size and nuclear staining for viable cells (DR-positive, DAPI-negative). For each sample, 50,000 to 200,000 cells were collected into 5 ml polystyrene tubes pre-filled with 4 ml AMB. Cells were pelleted at 350xg for 10 minutes in a swinging bucket centrifuge set at 6^°^C, cells resuspended in AMB at a concentration of 500 cells/µl and the cell suspension held on ice until microfluidic partitioning of single cells.

### Single Cell RNASeq and Data Analysis

An appropriate volume of each cell suspension estimated to contain 7,000 - 9,000 cells was combined with freshly prepared 10X Chromium reagent mix, and three zonal samples per replicate were loaded into separate lanes of a microfluidic partitioning device according to 10X Genomics Chromium v2 Single Cell 3’ reagent kit manufacturer’s instructions. Briefly, the Chromium v2 Single Cell 3’ protocol proceeded as follows: Cell capture, lysis and mRNA reverse transcription occurred in-droplets. cDNA recovered from the emulsion was cleaned-up, amplified by PCR, checked for size and yield on a 4200 Tape station (Agilent), processed into barcoded Illumina ready sequencing libraries, and again assayed for size and yield. The Translational Genomics Center at Children’s Hospital Los Angeles Center for Personalized Medicine carried out paired-end sequencing on the HiSeq 4000 platform (Illumina) using the HiSeq 3000/4000 SBS PE clustering kit (PE-410-001) and 150 cycle flow cell (FC-410-1002) and processed raw data into FASTQ files. Alignment of sequencing reads to the mouse (mm10) reference genome, as well as generation of BAM files and filtered gene-barcode matrices was accomplished by running Cell Ranger Single-Cell Software Suite 2.0 (10X Genomics) using the STAR aligner (Dobin et al., 2013) on the USC High Performance Cluster. The Cell Ranger *cellranger count* function output filtered gene-cell expression matrices removing cell barcodes not represented in cells. Principle component analysis and identification of variably expressed genes were carried out using the R packages Seurat (v 2.3.4) (Butler et al., 2018), ggplot2 (v. 3.0), Matrix (v.1.2-14) and dplyr (v 0.7.5) in R Studio. Briefly, for the primary analysis, the matrices from the 12 individual samples were first merged into a single metadata annotated r-object of 40,712 cells using the *Read10X, CreateSeuratObject AddMetaData* and *MergeSeurat* functions. The raw dataset was then filtered to remove genes expressed in less than three cells and the *FilterCells* function run with range filters for genes/cell (1,000-4,000) and mRNA transcripts/cell (UMIs; 1,000-16,000), and maximum percent mitochondrial genes/cell (35%); note renal epithelial cell types have high metabolic rates and expected higher mitochondrial gene detection than other cell types (Figure S1d, e, f). The resulting primary analysis dataset contained 19,125 genes across 31,265 cells. The *NormalizeData* function scaled and log transformed that dataset, from which 2464 variably expressed genes were identified with *FindVariableGenes*, followed by performing principle component analysis first on the variable genes with *RunPCA*, then extending to all dataset genes with *ProjectPCA*. The 30 useful principle components determined by the visualization tools *PCElbowPlot* and *JackStrawPlot* were used in graph-based clustering using *FindClusters* at resolution 1.0. *FindAllMarkers* generated the list of genes differentially expressed in each cluster compared to all other cells (Table S1) based on the Wilcoxon rank-sum test and limiting the analysis with a cutoff for minimum log FC difference (0.25) and minimum cells with expression (0.25). The secondary Seurat analyses of the nephron and ureteric epithelium (UE) profiles employed the *SubsetData* function to create new r-objects from cohorts of primary analysis clusters identified as nephron (4,6,8,10,12,13,18,25,26,27) or UE (5,7,20,23,17). Re-processing these with Seurat tools, as described above, identified contaminating non-type cells (vascular and immune) or unclassifiable profiles (stressed or mitotic) that were removed to yield the nephron dataset with 8,854 cells and the UE dataset with 4,349 cells. Visualizations using the *DotPlot* function and ggplot2 package graphically represent per cluster percentage expression (pct. exp.) by dot diameter and average nonzero expression in log2 scale (avg. exp.) by dot color; no dot shown if pct. exp. < 0.05.

### Analyses of cortical epithelial cell profiles at the nephron/ureteric epithelium junction

To analyze the transcriptional profiles of cortical epithelial cells at the junction between the nephron and UE at finer resolution, 1652 cells were computationally selected from the re-clustered nephron (2 clusters) and UE (5 clusters) datasets to create a new Seurat object. Reanalysis with the Seurat suite of tools yielded eight clusters with 18 to 551 cells [#0 551, #1 272, #2 258, #3 200, #4 198, #5 93, #6 62, #7 18]. Differentially expressed genes were identified using the *FindAllMarkers* function. The highly differentially expressed genes for each cluster were used in Pearson correlation analyses (run in R) to identify the most highly correlating genes and thereby identify candidate “marker genes.” Expression patterns were validated using the *FeaturePlot* and *DotPlot* functions.

### Lineage tracing of epithelium and marking ureteric epithelium

Six2TGC and Wnt4-GCE mice were generated as described previously (Kobayashi et al., 2008). For lineage tracing, Six2TGC male mice were crossed with homozygous *Rosa26^tdTomato^* females and their male offspring crossed with homozygous tgHoxb7-Venus females (Madisen et al., 2010; Chi et al., 2009). The Gdf15-CE has a nuclear GFP-F2A-CREERT2 cassette under control of the Gdf15 locus (unpublished strain McMahon laboratory). Rosa26^tdTomato^ mice (B6.Cg-Gt(ROSA)26Sortm14(CAG-tdT)Hze/J) were obtained from Jackson Laboratories and mated with the CRE-ERT2 lines. Tamoxifen was injected at E15.5 into pregnant Wnt4GCE/+, R26tdT/+ females or P2 Wnt4GCE/+, R26tdT/+ neonates and kidneys were collected at 8 weeks of age. Three experimental animals were sectioned and stained for this analysis. For Gdf15 lineage tracing experiments, the strain was crossed to R26tdT female mice and the resultant offspring injected with tamoxifen 3 days prior to analysis at adult stages.

### Immunohistochemical analyses of kidneys

Kidneys were harvested between 6-10 weeks from both male and female progeny. Three to four mice were used for each analysis. Mice were perfused with PBS followed by 4% PFA in PBS. Kidneys were then fixed for one hour in 4% PFA in PBS at 4°C. Kidneys were set in 4% low melting point agarose in PBS on ice and vibratome sectioned to a thickness of 200µm. Sections were collected into 6-well plates and kept suspended on a nutating platform during all washing and incubation steps. After blocking for one hour at 4°C in PBS with 2% SEA Block and 0.1% Triton X-100, primary antibodies were resuspended in the blocking solution and samples incubated at 4°C for 48 hours. To remove non-bound primary antibodies, the samples were washed for eight hours in five changes of PBS with 0.1% Triton X-100. Secondary antibodies diluted in blocking solution were applied at 4°C for 48 hours, followed by washing steps. For nuclear labeling, the samples were incubated at 4°C in 1 µg/ml Hoechst 33342 for two hours. Excess Hoechst was removed by PBS washes. Confocal imaging of vibratome slices (3D) was performed on a Leica SP8 using a 25x HC Fluotar L 25x/0.95 water immersion objective. The vibratome sections were imaged in 35mm MatTek glass bottom dishes. Vibratome images were opened and processed in LAS X (Leica), Imaris (Bitplane) and Photoshop (Adobe). Image brightness, contrast, and transparency were altered for 3D rending purposes to optimize resolution of cell distributions within the complex tissue. Immunofluorescent stains on cryo-sections were processed the same as above, except kidneys were embedded into OCT and 10µm sections generated (as described in below section for RNAscope).

### Antibody details

We used antibodies for immunofluorescent staining recognizing: Aqp1 (rabbit, Abcam, ab168387), Aqp2 (mouse, Santa Cruz, sc-515770), Atp6v1b1 (rabbit, Abcam, ab192612), Calb1(Calbindin-D-28K) (mouse, Sigma, C9848), Ckb (Mouse, MAB9076, R&D systems), GFP (chicken, Chemicon, AB16901), Hoxd10 (rabbit, Abcam, ab76897), Jag1 (goat, R&D Systems, AF599), Slc26a4 (rabbit, Sigma, SAB2104723), Slc4a1 (rabbit, Alpha Diagnostic, AE11-A1), Slc5a2 (rabbit, Abcam, ab85626), Rhbg (rabbit, Novus, NBP2-33527), and Tfcp2l1 (goat, R&D Systems, AF5726). Complete information is provided in tabular form in Table S6.

### RNAscope details

RNAscope probes were obtained from Advanced Cell Diagnostics. *In situ* hybridization was performed following RNAscope Multiplex Fluorescent Reagent Kit v2 user manual (document #323100-USM). Briefly, perfusion with PBS, kidneys were harvested and fixed at 4°C in 4% PFA in PBS for 1 hour, followed by overnight at 4°C in 30% sucrose. Kidneys then were embedded in OCT in molds floated on a dry ice/ethanol bath. Cryosections (10 µm) were fixed in 4% PFA at 4°C overnight before the standard pretreatment steps for fixed frozen tissue sample in the protocol. Signal amplification after the RNAscope Multiplex fluorescent v2 assay was performed with TSA plus fluorophores as recommended and Alexa Fluor conjugated secondary antibodies (Thermo Fisher Scientific). Complete details of RNAscope probes used here are provided in Table S6.

### qPCR

Total RNA was isolated from FACS sorted cells or homogenized kidney tissues using RNeasy mini kits (Qiagen), followed by cDNA synthesis with Superscript Vilo kit (Thermo Fisher Scientific). qPCR was performed with SYBR green on Applied Biosystems 7500 fast real-time PCR system. Reported Ct values are the mean of three technical replicates on the same cDNA prep. All PCR assays reported are sample size (n) = 1. The details of primers used are presented in Table S6.

### Gene Ontology

Gene Ontology analyses were carried out in PANTHER (Mi et al., 2013). For analyses of cortical epithelial profiles around the N/UE junction, the top 50 differentially expressed genes from relevant clusters were used to identify biological processes. For analyses of sex-biased PT profiles, Seurat *FindMarkers* function was run to identify differentially expressed genes that distinguish a cluster from specific other clusters. The four comparisons shown in Figure S2 were for f S3: 9 vs 9m; m S3: 9m vs 9; f PT: 0,3,9 vs 2,5,9m and m PT: 2,5,9m vs 0,3,9.

### Website

Kidney Cell Explorer (https://cello.shinyapps.io/kidneycellexplorer/) was developed using Shiny (Chang et al., 2018)and the R programming language. We utilized interactive visualization modules of the VisCello package (Packer et al., 2019) and adapted them to allow exploration of kidney specific data. Key features of the website include interactive t-SNE plots, feature expression plots, metacell expression heatmap and detailed ontology annotation Expression heatmaps on the webpage show gene expression pattern across nephron and ureteric epithelium cells stratified by ontology based “metacells.” We assigned Seurat clusters to metacells associated with kidney ontology based on enrichment of curated signature genes as described in the main text. We then computed average Seurat normalized gene expression and proportion of non-zero expression cells for each gene across metacells. The number of genes recovered at metacell level is significantly higher than the detected gene number per single cell, allowing more robust and comprehensive profiling of positional expression patterns.

## Supplemental Information

**Figure S1.**
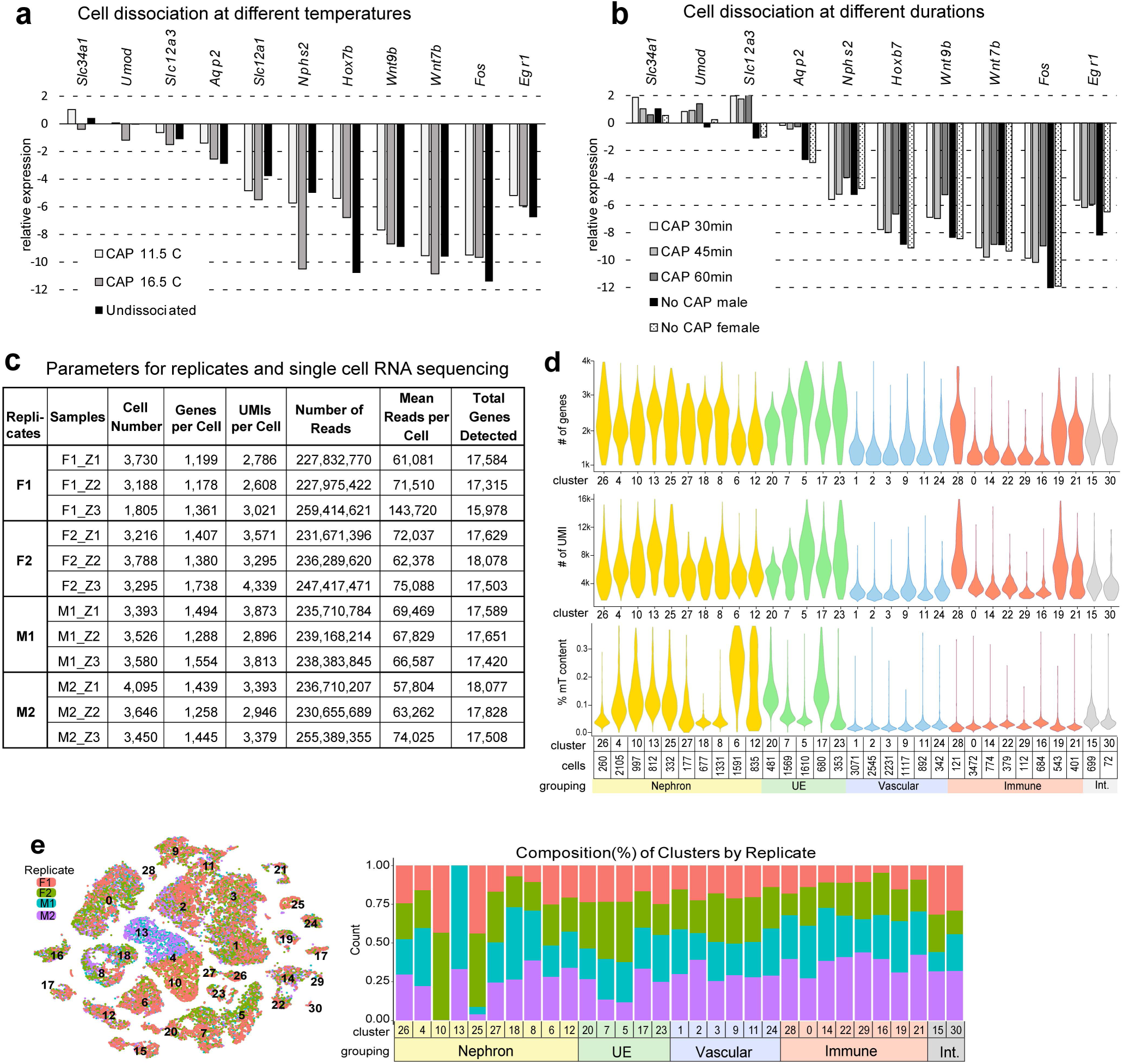
Optimizing dissociation and primary analysis metrics *Related to Figure 1*. **a-b**, Plot of qPCR results shown as dCt ([Ct_Gapdh]-[Ct_gene-of-interest], *Gapdh* set to zero) comparing cDNA yields for representative cell type enriched gene markers and stress response genes (*Egr1, Fos*) from cold active protease (CAP) dissociation varying temperature (a) and time (b) of dissociation. ‘Undissociated’ and ‘No CAP’ samples were directly extracted adult kidney samples. **c,** Sequencing metrics reported by Cell Ranger for all 12 primary tissue samples (4 kidneys × 3 zones). **d**, Primary analysis dataset graphically represented as violin plots of all 30 clusters (ordered as in Figure 1c) showing per cluster levels for nGenes, nUMI and percent mitochondrial genes. Notably, distal tubule cells (clusters 6,12), have high percent mitochondrial genes while showing typical RNA complexity (nUMI), suggesting high metabolic rates in good quality cells. **e,** tSNE and stacked barplots of the primary dataset (as in Figure 1b-d), illustrating the distribution (left) and composition (right) of the clusters with respect to the four replicates. F: female, M: male, UE: ureteric epithelium, Int: interstitial.

**Figure S2.**
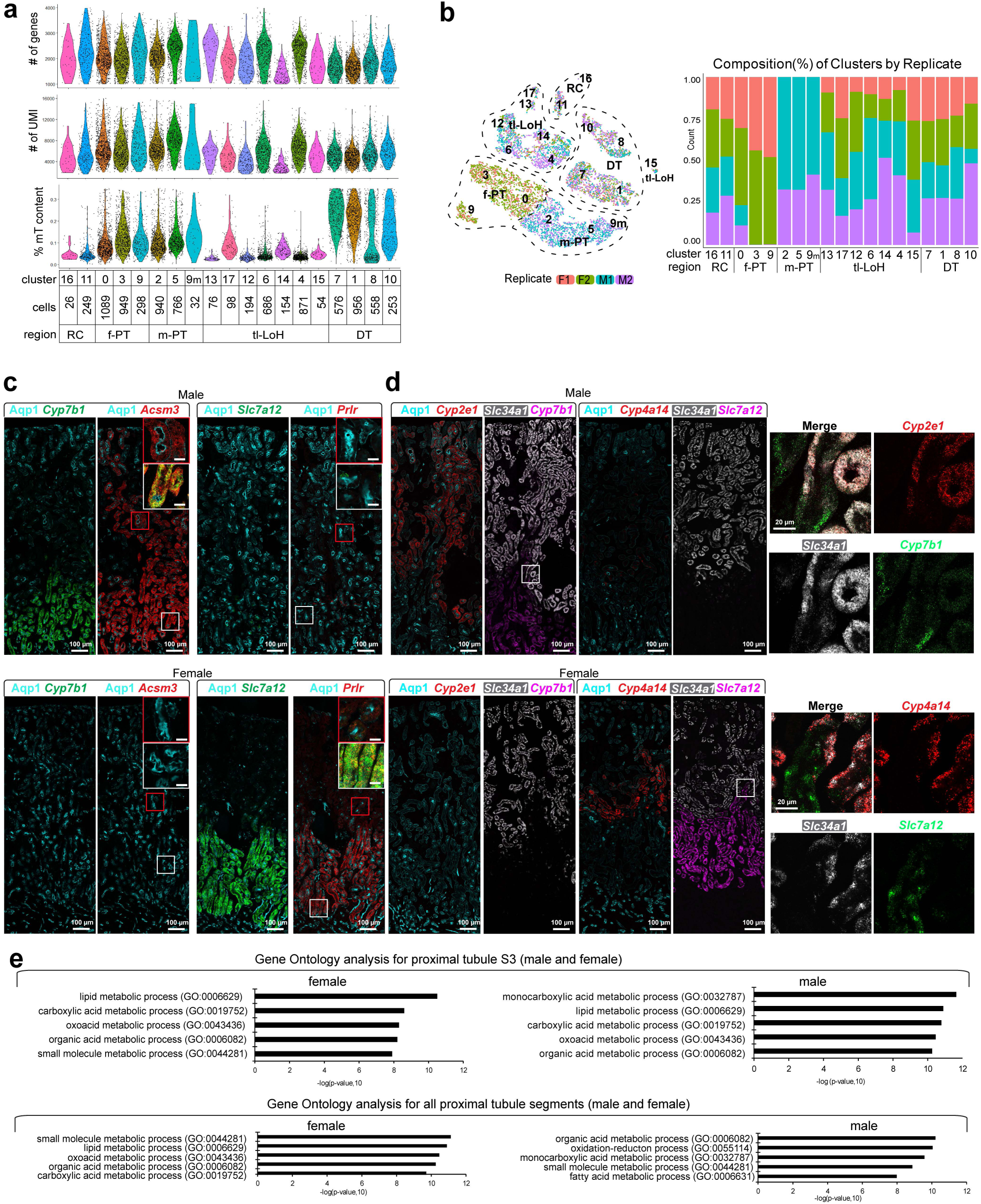
Nephron dataset metrics and sex bias validation for the proximal tubule *Related to Figure 2*. **a**, N dataset as violin plots of the 19 clusters in Figure 2b, showing per cluster levels for nGenes (top), nUMI (middle) and percent mitochondrial genes (bottom). **b**, tSNE and stacked barplots of N dataset illustrating the distribution (left) and composition (right) of the clusters by replicate. **c**, Expression of sex biased, PT S3+S2 enriched markers *Ascm3* (m) and *Prlr* (f) in adult kidney tissue by RNAscope *in situ* and antibody labelling; inset scale bar 20µm. **d**, Expression of segment restricted, sex biased PT S2 markers are non-overlapping and demarcate the boundary between PT S2 and S3 segments. Scales and probes indicated on each panel. **e-f**, Gene Ontology analyses showing the top five GO Terms returned for genes showing a sex specific enrichment in female or male S3 (e) or throughout all female and male PT segments (f). RC: renal corpuscle, f-PT: female proximal tubule, m-PT: male proximal tubule, tl-LoH, Loop of Henle-thin limbs, DT: distal tubule.

**Figure S3.**
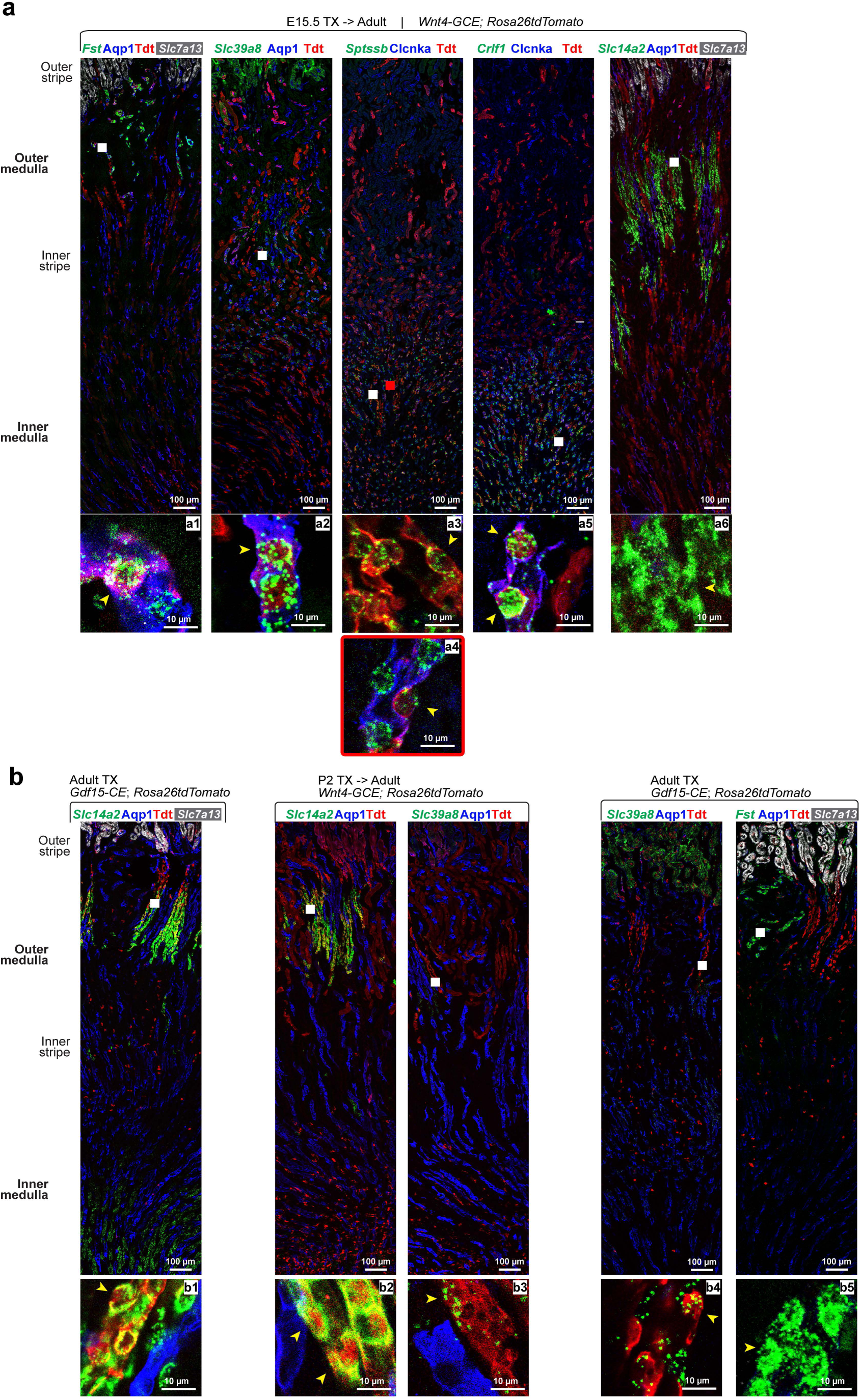
Localization of thin Loop of Henle markers in early vs late forming nephrons *Related to Figure 3*. **a-b**, Low power views of thin limbs of Loop of Henle (tl-LoH) markers in tdT-labelled adult kidney sections; boxes indicate region of inset below each panel. Boxes also show where high power images in Figure 3e,f were collected relative to medullary depth shown at left; text above field details genetic combinations and tamoxifen (TX) injections; scale bar and probes indicated on each panel; arrowheads: a1, *Fst*^+^/Apq1^+^/tdT^+^; a2, *Slc39a8*^+^/Apq1^+^/tdT^+^; a3, *Sptssb*^+^/Clcnka^-^/tdT^+^; a4, *Sptssb*^+^/Clcnka^+^/tdT^+^; a5, *Crlf1*^+^/Clcnka^+^/tdT^+^; a6, *Slc14a2*^+^/Apq1^-^/tdT^-^; b1-2, *Slc14a2*^+^/Apq1^-^/tdT^+^; b3-4 *Slc39a8*^+^/Apq1^-^/tdT^+^; b5, *Fst*^+^/Apq1^-^/tdT^-^.

**Figure S4.**
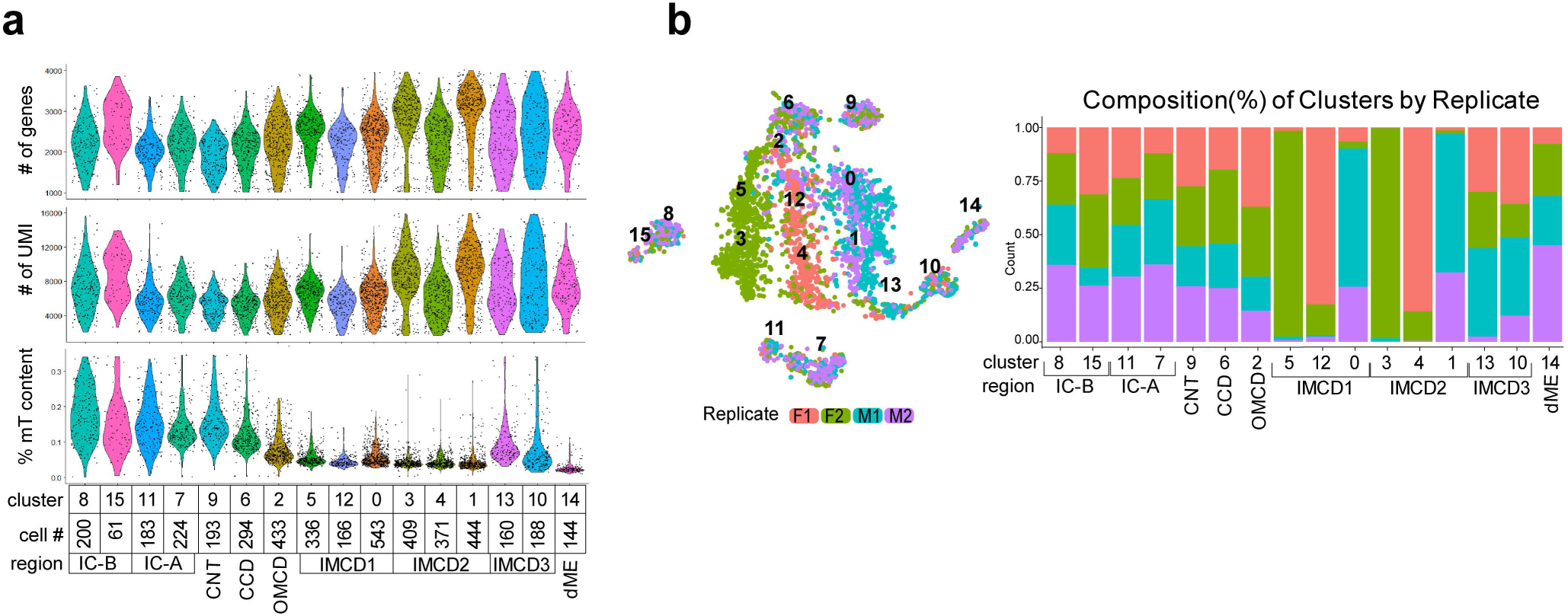
Dataset metrics for ureteric epithelial cell groupings *Related to Figure 4*. **a**, UE dataset as violin plots of the 16 clusters in Figure 4b, showing per cluster levels for nGenes (top), nUMI (middle) and percent mitochondrial genes (bottom). **b**, tSNE and stacked barplots of UE dataset illustrating the distribution (left) and composition (right) of the clusters by replicate; IC, intercalated cells, type A and B; CNT, connecting tubule; CCD, cortical collecting duct; OMCD, IMCD, outer and inner medullary collecting duct, types1-3; dME, deep medullary epithelium.

**Figure S5.**
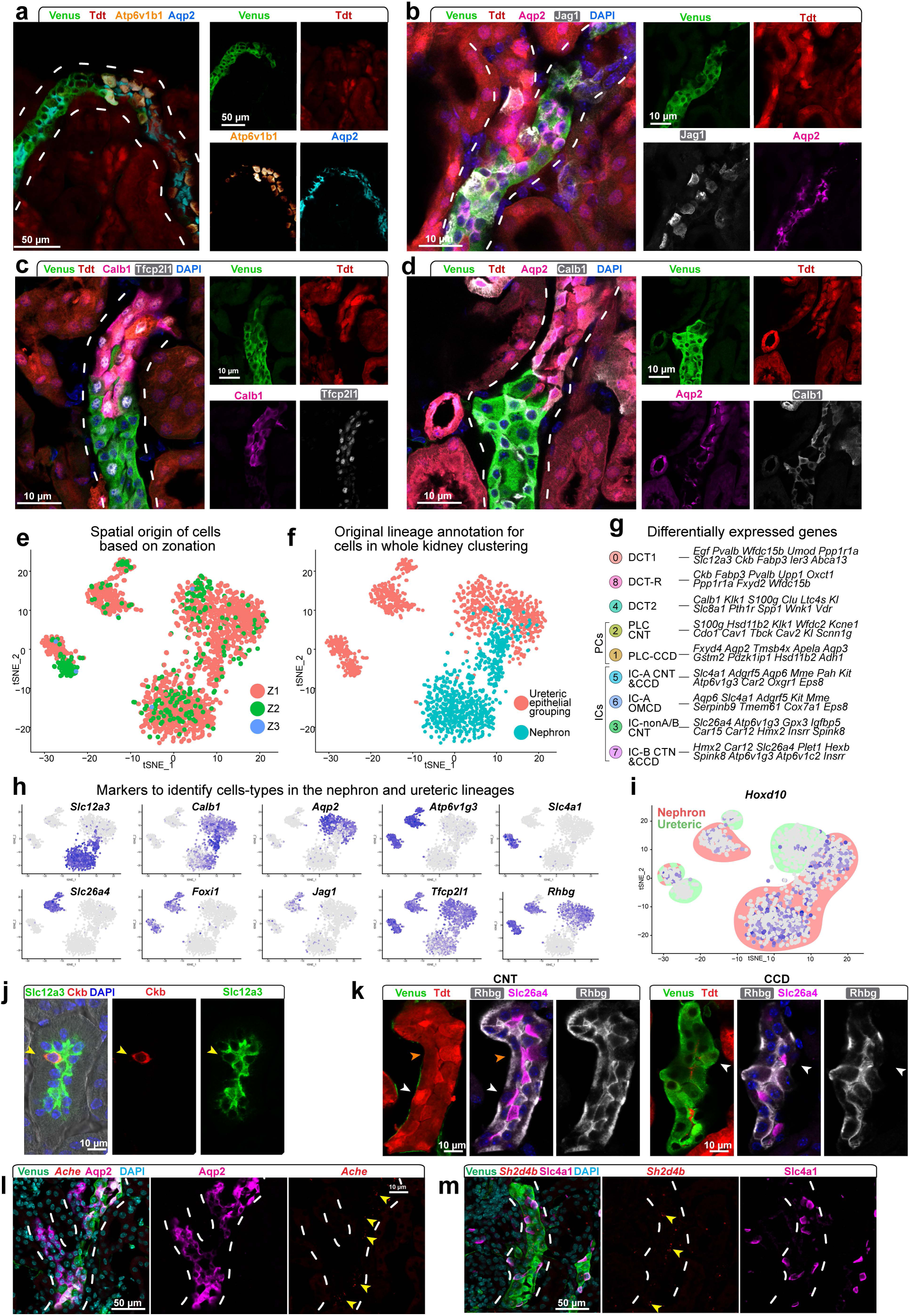
Nephron lineage derived principal and intercalated cell types at connection to collecting system *Related to Figure 5*. **a**, General cell markers of intercalated cells (IC, Atp6v1b1) and principal cells (PC, Aqp2) in nephron (N) and ureteric epithelium (UE) around the junction with collecting system. **b-d**, Cell specific analysis of indicated markers in PC, IC and other epithelial cell types around the junction with collecting system. **e,f**, tSNE projections of re-clustered cortical epithelial cell types from N and UE datasets displayed as zone (e) or lineage (f) of origin. **g**, Differentially expressed genes for each cluster in Figure 5i. **h**, Feature plots showing expression of key marker genes defining cell types. **i**, Feature plot displaying *Hoxd10* expression across clusters in Figure 5i. **j**, Rare Ckb^+^ cells (arrowhead) within the Slc12a3^+^ distal convoluted tubule cluster 8 in Figure 5i. **k**, NonA-nonB IC identified by double label for Rhbg and Slc26a4; arrowheads: white, IC-B; orange, nonA-nonB IC. **l-m**, Immunofluorescent and RNAscope validation (arrowheads) of Ache in UE-PCs and *Sh2d4b* in UE-IC-B, respectively; merged panel on left.

**Figure S6.**
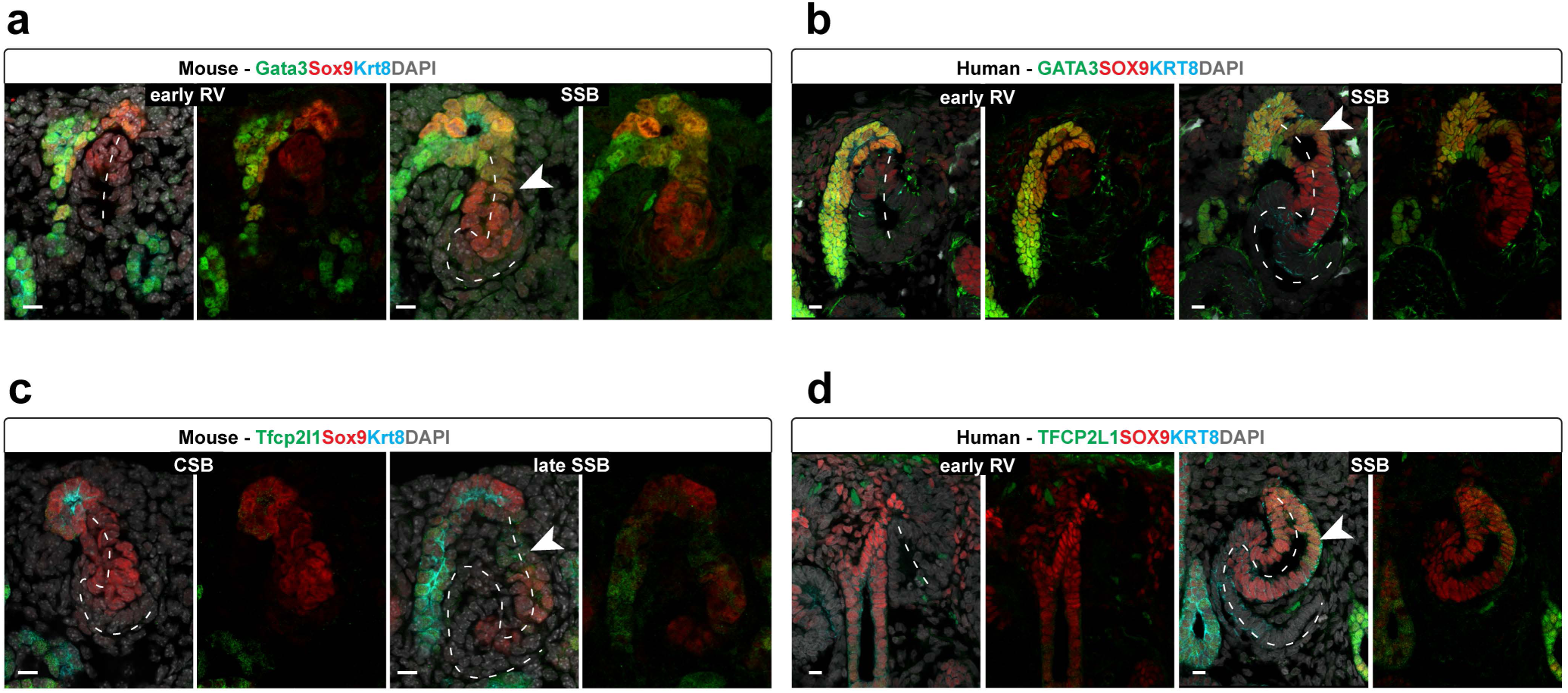
Progressive shift of transcription factors from the ureteric lineage into the distal nephrogenic lineage in early nephrogenesis. **a-d**, Human (week 15) and mouse (day of birth) kidney sections immunostained to examine cellular distributions of transcription factors first expressed in the UE and subsequently in distal cell types of the developing nephron. Nephron development initiates with the epithelial renal vesicle (RV), proceeds though a Comma-shaped body (CSB) to an S-shaped body (SSB). A patent luminal interconnection is generated between the nephron and collecting system by the SSB stage; dashes follow proximal-distal axis of nephron precursors; arrowheads indicate transcription factors detected in distal nephron. Scale bar 10µm.

**Figure S7.**
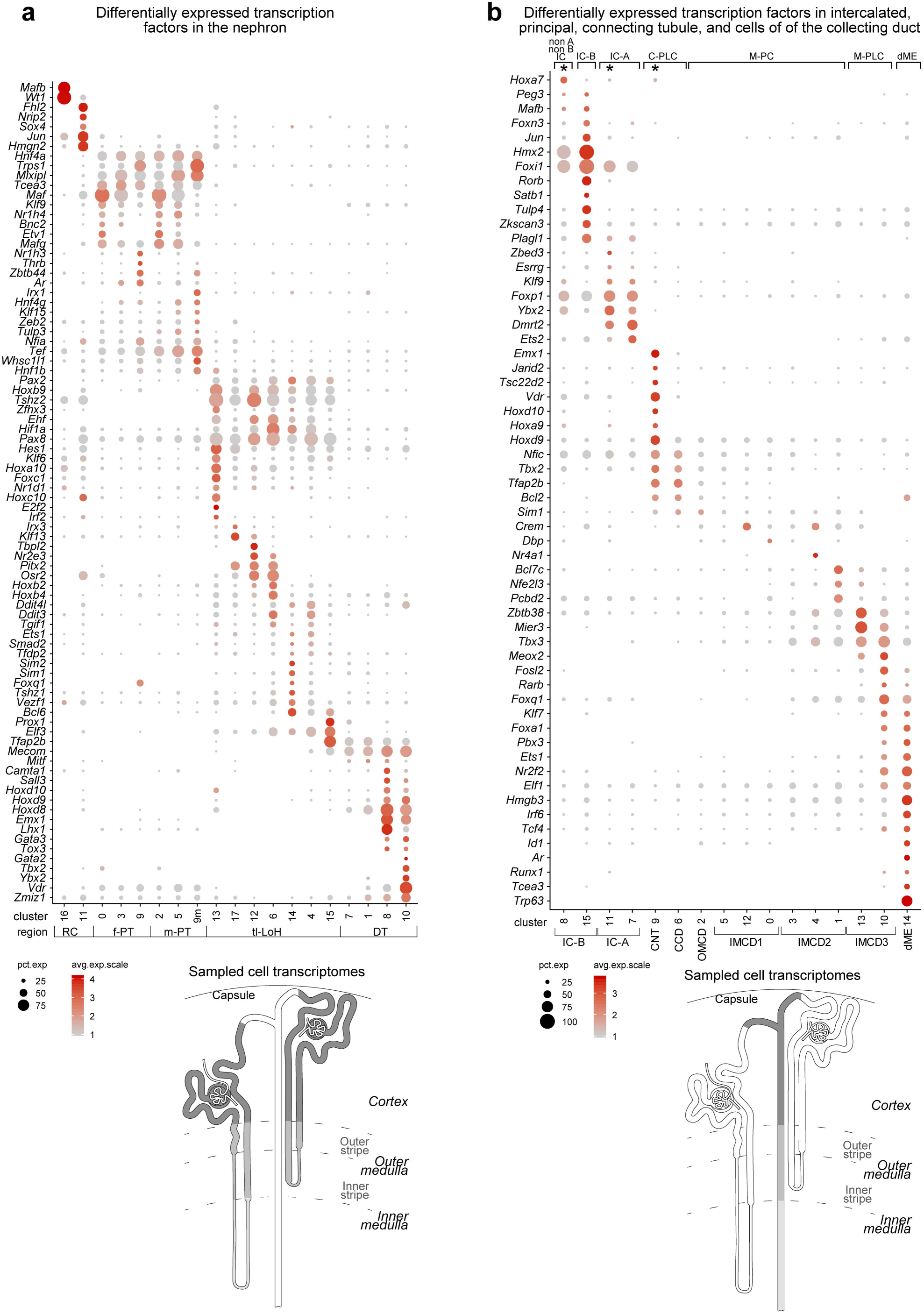
Enriched expression of transcription factors in clusters of nephron and ureteric epithelium datasets. **a-b**, Dot plots showing differential expression of transcription factors along the proximal-distal axis of the mature nephron (a) and ureteric epithelium of the collecting system (b), with a schematic (below) indicating the region sampled: RC, renal corpuscle; f-PT, female proximal tubule; m-PT, male proximal tubule; tl-LoH, thin limbs of Loop of Henle; DT, distal tubule; IC, intercalated cell, type a and b; CNT, connecting tubule; CCD, cortical collecting duct; OMCD, outer medullary collecting duct; IMCD, inner medullary collecting duct; dME, deep medullary epithelium.

**Figure S8.**
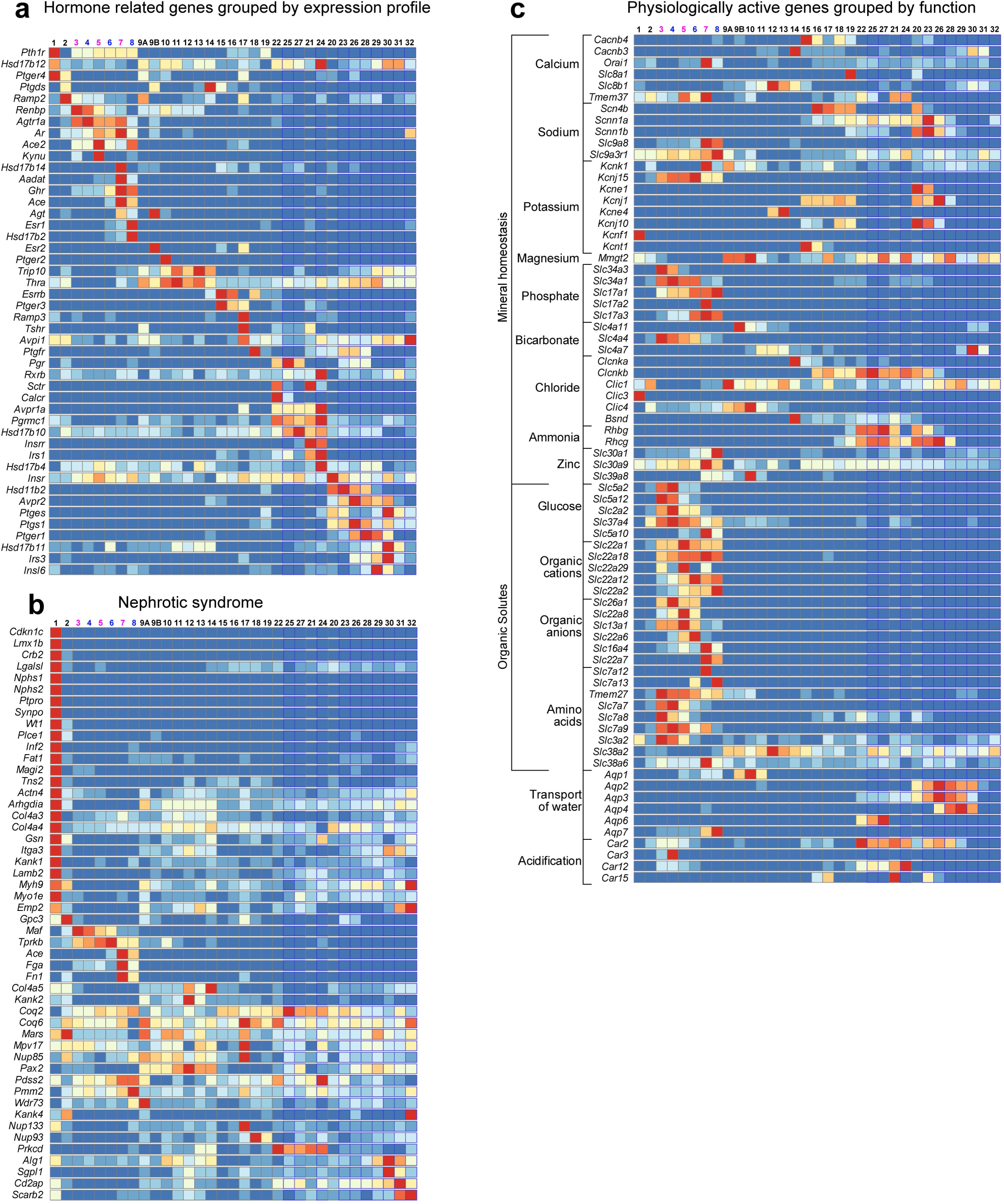
Kidney Cell Explorer batch search outputs heatmaps spanning all metacells. **a-c**, Heat maps of gene expression for multi-gene searches in metacells of nephron and collecting duct showing examples for batch searches with genes linked to hormone regulation (a), nephrotic syndrome (b) and specific physiological activities (c).

## Supplemental Tables

**Table S1** Mouse adult kidney dataset lists of top 50 most differentially expressed genes for each cluster shown in the tSNE projection in Figure 1b. See tab entitled “lineage table” for key to content, Excel file. *Related to Figure 1 and Figure S1*

**Table S2** Nephron dataset lists of differentially expressed markers in each cluster shown in the tSNE projection in Figure 2b. See tab entitled “read me” for key to content, Excel file. *Related to Figure 2 and Figure S2*

**Table S3** Details of Gene Ontology analyses for genes expressed with sex bias in proximal tubules as shown in Figure S2e for entire proximal tubule and S3 regions. Separate tabs for PT gene lists used and Panther results tables, Excel file. *Related to Figure S2*

**Table S4** Ureteric epithelium dataset lists of differentially expressed markers in each cluster shown in the tSNE projection in Figure 4b. See tab entitled “read me” for key to content, Excel file. *Related to Figure 4 and Figure S4*

**Table S5** Combined list of differentially expressed markers in the combined cortical clusters dataset shown in the tSNE projection in Figure 5i, Excel file. *Related to Figure 51 and Figure S5*

**Table S6** Reagent usage table with separate tabs for Antibodies, RNAscope probes and PCR primers, Excel file.

